# *De novo* design of triosephosphate isomerases using generative language models

**DOI:** 10.1101/2024.11.10.622869

**Authors:** Sergio Romero-Romero, Alexander E. Braun, Timo Kossendey, Noelia Ferruz, Steffen Schmidt, Birte Höcker

## Abstract

The design of proteins with tailored functions is of immense interest to biotechnology, medicine, and the chemical industry. While protein design is rapidly evolving with the use of AI techniques, the design of complex enzymes remains a challenge. Here, we present the use of two large language models (LLMs), ZymCTRL and ProtGPT2, for the generation of de novo enzymes that catalyze the triosephosphate isomerase (TIM) reaction. Natural TIM enzymes are obligatory oligomers that catalyze a multi-step isomerization reaction near the diffusion limit. This makes TIM an ideal target to assess the generative ability of protein language models. Newly generated sequences were filtered to obtain a set of twelve candidates from each approach for experimental validation. Multiple constructs from both language models exhibit the intended function in vivo through their ability to complement a TIM-deficient E. coli strain. In-depth characterization of the best-behaving artificial enzyme reveals behavior and catalytic efficiency close to its natural counterparts. These findings support the use of conditional and fine-tuned unconditional LLMs for the generation of complex enzymes.

## Introduction

The design of tailor-made enzymes has been a long-standing goal in biochemistry. With powerful computational algorithms, it has become possible to generate proteins from scratch that fold into predefined geometries (Pan and Kortemme 2021; Korendovych and DeGrado 2020; Huang et al. 2016), build high-affinity drug binders (Polizzi and DeGrado 2020; Vázquez Torres et al. 2024), create proteins and antibodies that are bioactive *in vivo* (Rhys et al. 2022; Herud-Sikimić et al. 2021; Sesterhenn et al. 2020; Donnelly et al. 2018), and introduce basic catalytic sites into existing proteins (Korendovych et al. 2011; Rajagopalan et al. 2014; Villali and Kern 2010; Moroz et al. 2015; Privett et al. 2012; Siegel et al. 2010; Bjelic et al. 2013). However, building small molecule binding sites and sites for more complex enzymatic reactions remains a challenge due to necessary precision in residue placement and balancing compact structure with enough flexibility for a function to efficiently occur (Lechner et al. 2018; Villali and Kern 2010). The recent advent of AI-based methods for structure prediction (Jumper et al. 2021; Lin et al. 2023; Baek et al. 2021) has had a strong impact on protein design and is changing the field rapidly (Khakzad et al. 2023). Inspired by these successes, a range of new tools, such as ProteinMPNN or RFdiffusion (Dauparas et al. 2022; Watson et al. 2023), have been developed for the design of new sequences and structures.

Apart from these structure-centered approaches, it is attractive to explore the ability of natural language processing (NLP) tools for the design of novel proteins directly from sequence. Recent advances in NLP have been paralleled by new developments in biochemistry, with research tools emerging concomitantly with NLP innovations (Ferruz and Höcker 2022). Within this field, the use of large language models (LLMs) for protein sequence embedding, annotation, and generation has been shown to be an effective method for exploring uncharted protein sequence space (Schmirler et al. 2024; Romero-Romero et al. 2023). Among these models, ProtGPT2 is an unconditional model able to generate novel protein sequences with diverse non-idealized structures, topologies and sizes (Ferruz et al. 2022). Also on the tenet of sequence generation, the conditional protein language model ProGen was built and applied to five different lysozyme families after extensive fine-tuning (Madani et al. 2023). The model ZymCTRL, which is based on the CTRL transformer similar to ProGen, was developed for the conditional generation of putative enzymes according to user-input based on the EC classification (Munsamy et al. 2024). The evolutionary scale model ESM2 combines LLMs with other conditional models, following a fixed backbone and an unconstrained generation approach (Verkuil et al. 2022).

Here, we explore the ability of two different LLMs to generate functional enzymes that catalyze the complex triosephosphate isomerase (TIM) reaction with sequences distant to the natural enzymes. TIM (EC 5.3.1.1) is the eponymous enzyme of the TIM-barrel fold (Alber et al. 1981; Nagano et al. 2002; Banner et al. 1975) that catalyzes the reversible interconversion of glyceraldehyde-3-phosphate (GAP) and dihydroxyacetone phosphate (DHAP), with a catalytic efficiency close to the diffusion limit (Albery and Knowles 1976; Go et al. 2010; Knowles 1991). TIM is involved in multiple metabolic pathways and its proper function is crucial for the energetic balance in cells (Orosz et al. 2006). Its topology is characterized by the ubiquitous (ꞵ/ɑ)_8_-barrel (Romero-Romero et al. 2021) with a catalytic triad located at the C-terminal face of the ꞵ-strands consisting of a lysine (providing electrostatic stabilization of the bound substrate), a histidine (electrophilic catalyst), and a glutamic acid (catalytic base) (Richard 2012). This essential metabolic enzyme is only active as a homooligomer (dimer or tetramer) in nature since the correct positioning of the active site is facilitated by the insertion of the oligomerization loop from the neighboring monomer. The obligatory oligomerization, the high activity in natural members of this enzyme class, and the defined topology make this enzyme a suitable model to assess the performance of state-of-the-art protein language models (pLMs) in recapitulating the complex sequence-structure-function relationships inherent in proteins.

Here, we tested the ability of two different sequence generation approaches using the transformer-based pLMs ZymCTRL and ProtGPT2 to create novel proteins that catalyze the desired TIM reaction. Success was investigated experimentally by *in vivo* complementation and *in vitro* enzyme assays. We discuss the importance of a well-curated dataset for training, show the effect of filtering on sequence performance, and comment on the ability of pLMs to recapitulate interactions throughout the entire sequence. Overall, our work highlights the use of pLMs as a new viable and robust method for enzyme design generating highly diverse sequences in an enzyme class without using large-scale screening methods.

## Results and Discussion

### ZymCTRL generates triosephosphate isomerase sequences with nature-like properties

Autoregressive sequence generation can be sorted into two categories: conditional and unconditional generation (Romero-Romero et al. 2023). Conditional models are trained with tags coupled to the respective training-input and therefore able to generate output that recreates the essence of the given tag used as a prompt. Unconditional models are trained without information coupled to a dataset resulting in unbiased generation. ProtGPT2 (Ferruz et al. 2022) and ZymCTRL (Munsamy et al. 2024) are representatives of unconditional and conditional pLMs, respectively. ZymCTRL was trained on the enzyme subset of the UniProt database where each sequence was coupled to the corresponding activity represented as the enzyme commission (EC) number. The coupled training allows for conditional generation of putative enzymes affiliated with the EC number as input term. ProtGPT2, on the other hand, was trained on the UniRef50 database containing proteins throughout all classifications and folds without any conditioning. Because of the more defined generation process, the conditional generation of TIMs was explored first (Figure 1).

**Figure 1.**
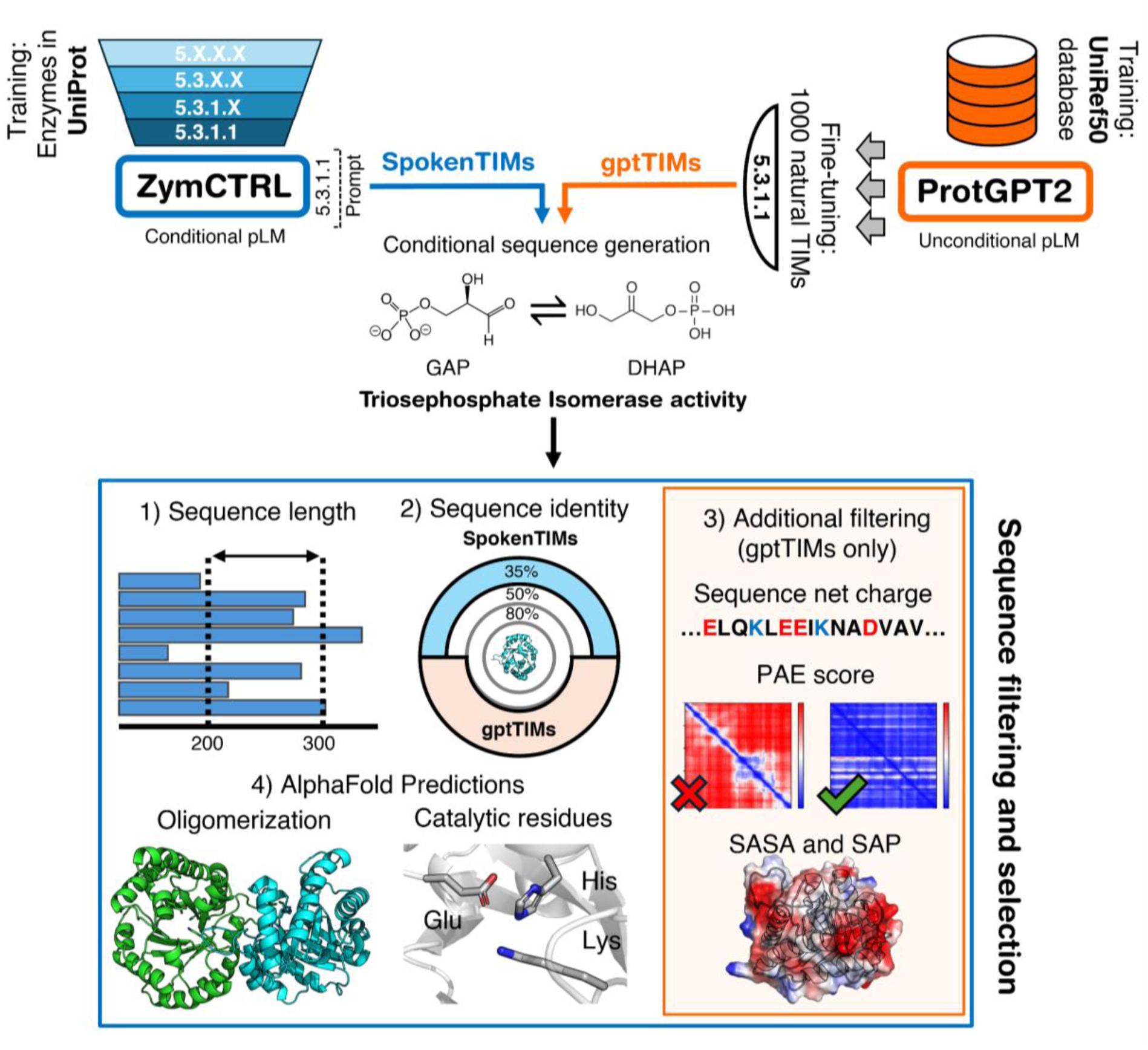
Generation and filtering of novel TIM sequences using a conditional (ZymCTRL: SpokenTIMs) and an unconditional (ProtGPT2: gptTIMs) language model. Triosephosphate isomerase sequences were generated with ZymCTRL, using the EC number 5.3.1.1 that represents the TIM reaction as input. For the generation of sequences using ProtGPT2, the model was fine-tuned with ∼1,000 sequences classified as TIMs in UniProt. Sequences with a length between 200 and 300 amino acids, a typical range for natural TIMs, were selected (**1**). ZymCTRL generated sequences (SpokenTIMs) with a sequence identity to any natural protein > 35 % were discarded, while for ProtGPT2 generated sequences (gptTIMs) the cut-off was set to 50 % (**2**). gptTIMs were further filtered by their net sequence charge, average PAE value, and hydrophobic surface parameters (SASA and SAP scores) of the predicted ColabFold models (**3**). Passing sequences of both pLMs were modeled as oligomers using ColabFold and checked for existence and proper predicted orientation of the catalytic residues (**4**). pLM: protein language model, GAP: Glyceraldehyde 3-phosphate, DHAP: Dihydroxyacetone phosphate.

Via ZymCTRL, we generated about 90,000 sequences using the EC number 5.3.1.1 as input prompt and the pre-trained model, i.e. without further fine-tuning the model in any additional, curated dataset. Despite no additional restraints, the resulting sequences mimicked the length distribution of the EC 5.3.1.1 dataset (Figure S1A). To counteract the influence of misannotated TIM sequences in the training data, the generated sequences were filtered to be between 200 to 300 amino acids in length corresponding to the average length of natural TIMs (247 ± 57 amino acids). 93.4 % of the sequences were located in this range. The majority of these filtered sequences are diverse and share a sequence identity of 50 % or less (Figure S2). The distribution of sequence identities corresponds to the distribution among natural TIMs. These findings highlight the influence and importance of the training dataset on sequence generation and display that pLMs recapitulate a broad variety of properties from the protein primary structure.

Concerning sequence identity to natural proteins, the generated sequences show major populations at 55 % and 90 % identity (Figure S3). Sequences with identities >35 % to any natural protein were discarded to assess the ability of the pLM to generate TIMs distant from natural ones and closer to the twilight zone (Table S3). ColabFold models of the 87 remaining putative enzymes were evaluated based on their propensity to adopt the (βα)_8_-barrel fold and the existence and positioning of the three catalytic residues Lys, His, and Glu. We further checked if the candidates are predicted to be either homodimers or homotetramers (Table S3) since this enzyme is only active as a homooligomer (Olivares-Illana et al. 2017). The twelve most promising sequences, named SpokenTIMs, were selected and experimentally characterized. All twelve are classified as TIMs by CATH (Sillitoe et al. 2021), SCOPe (Fox et al. 2014), HHPred (Zimmermann et al. 2018; Soding et al. 2005), and Foldseek (Van Kempen et al. 2023) and share a maximum sequence identity of 35% with each other (Figure S5).

### Language-model-designed TIMs complement a TIM-deficient E. coli strain

TIM is a key enzyme in several metabolic pathways and of utmost importance for the energy balance of the cell (Schümperli et al. 2007). Hence, activity of SpokenTIMs was assayed *in vivo* using an *E. coli* strain lacking the natural gene for TIM (Saab-Rincón et al. 2001). Selection can be performed using either glucose or glycerol as a sole carbon source since the lack of TIM in the cell results in low to no growth, with glycerol providing the most stringent selection where growth rate is completely dependent on TIM activity. In complementation assays, the cells containing a SpokenTIM variant that harbors TIM activity will show increased growth based on the level of activity and solubility of the respective protein.

From the twelve selected SpokenTIMs, three led to complementation. In particular, SpokenTIM9 showed a clear growth on plates containing either carbon source after 48 hours compared to 24 hours for three natural TIMs that represent all clades of life (Figure 2A and Figure S6). Further, SpokenTIM4 and SpokenTIM5 show low growth after the same time. Based on this assay three out of twelve ZymCTRL-designed enzymes are functional *de novo* TIMs. The rapid growth of TIM-deficient cells carrying the SpokenTIM9 gene indicates that this construct is the most active and/or soluble of all SpokenTIMs.

**Figure 2.**
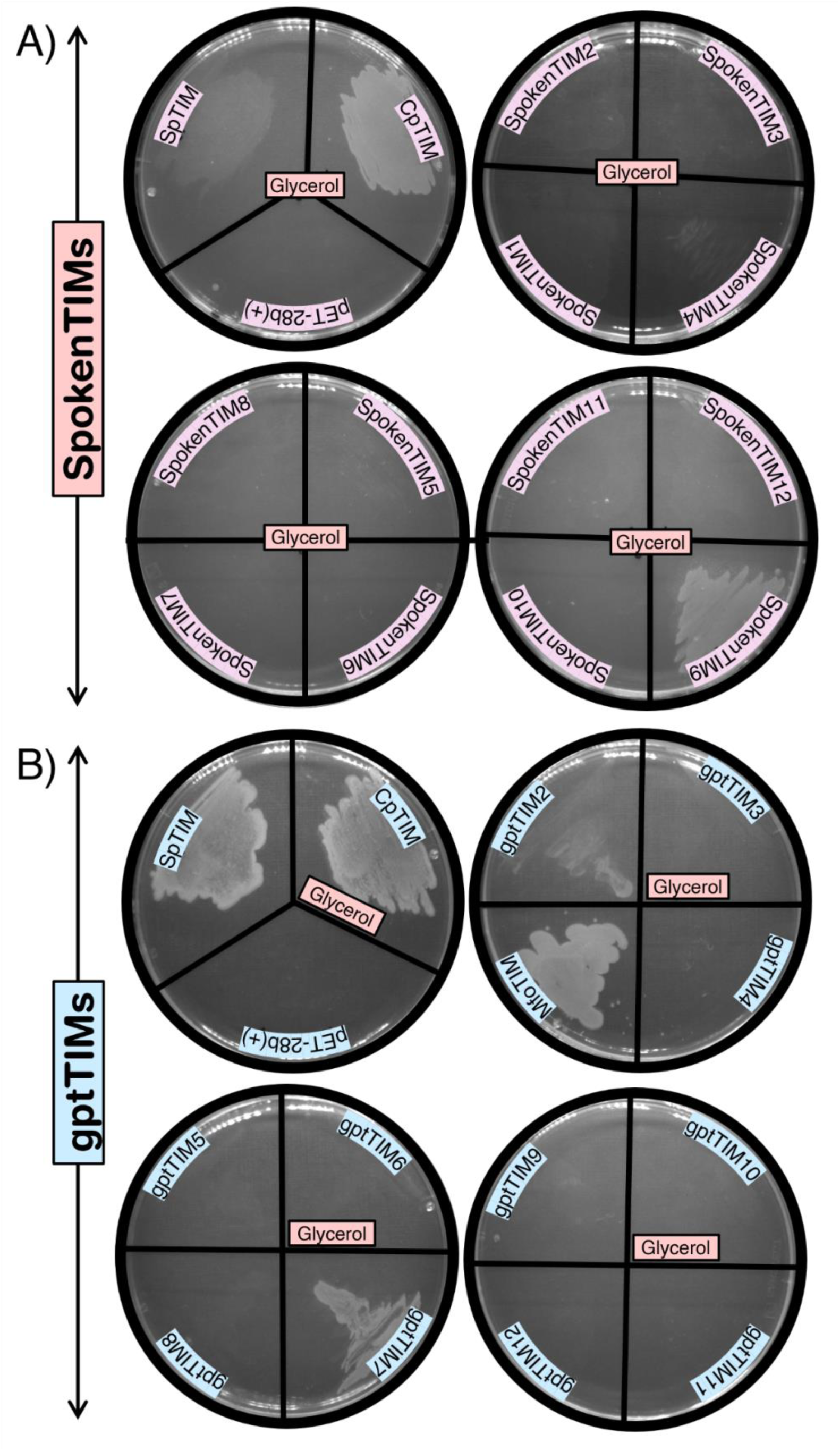
SpokenTIMs and gptTIMs can complement triosephosphate isomerase deficiency in an *E. coli* TIM deletion strain. *E. coli* BL21 cells lacking the native gene for triosephosphate isomerase, but carrying the SpokenTIM or gptTIM constructs, were plated on M9 minimal media containing glycerol as the sole carbon source. (**A**) Snapshot after 48 hours: SpokenTIM9 activity complements the gene-deficiency resulting in increased growth rates in comparison to cells carrying inactive/insoluble proteins or an empty pET28b(+) vector. Also SpokenTIM4 and SpokenTIM5 show slight growth. (**B**) gptTIM2 and 7 also show increased growth rates on glycerol after 48 hours compared to cells carrying inactive proteins or an empty pET28b(+) vector. As positive controls cells carrying the natural TIMs from *Clostridium perfringens, Schizosaccharomyces pombe,* or *Methanotorris formicicus* were used.

### Fine-tuned ProtGPT2 enables generation of functional TIMs

Based on the performance of SpokenTIM9, this sequence was used to reevaluate our filtering pipeline for pLM generated sequences. So far, the quality of sequences was assessed simply by sequence length constraints, visual inspection of predicted structures, and the presence and predicted positioning of the three residues necessary to catalyze the TIM reaction. We decided to evaluate further criteria. As suggested by Verkuil et al. (Verkuil et al. 2022) we removed all sequences with a net charge between 2 and −2 as low protein charges promote aggregation (Ventura 2005). It is known that soluble proteins require predominantly hydrophilic amino acids on their surface. Therefore, low spatial aggregation propensity (SAP) scores are indicative of soluble proteins, as exposed hydrophobic patches on a protein surface contribute to the total SAP score. Predicted models of the generated proteins with average SAP scores over all residues lower than 0.4 were selected for further filtering. In relation to this metric, the hydrophobic solvent accessible surface area (SASA) was determined, as a highly hydrophobic surface area suggests a protein to be prone to aggregation. We excluded all proteins with a hydrophobic surface area greater than 1.7 times the ideal surface area of a globular protein (Dill et al. 2011). Additionally, the PAE score of the ColabFold prediction was utilized as a filtering metric. Monomers with average PAE scores lower than 5 Å were observed to be well predicted, as well as having a high probability to fold into the desired structure (Watson et al. 2023).

The filtering was applied to the putative TIMs generated by ZymCTRL. Interestingly only SpokenTIM9 passed this filtering. This construct being the best performing of the twelve experimentally characterized SpokenTIMs, supports the idea that the filtering can guide sequence selection for experimental characterization.

To test this hypothesis and explore the capabilities of another type of directed generation of proteins with desired attributes (Figure 1), the generative protein language model ProtGPT2 was fine-tuned on natural TIM sequences. To retrieve sequences distant to known proteins, the generated sequences were filtered to be less than 50 % identical to proteins in the nr database, where 32.6% of these artificial sequences aligned to natural TIMs. The sequences were further filtered as previously described. Simply filtering by physicochemical properties and ColabFold quality parameters resulted in 89.9 % of the remaining sequences aligning to natural TIMs. This indicates that the fine-tuning of the unconditional model ProtGPT2 drives the generation towards sequences of wanted properties. We observed that filtering for favorable protein attributes further enriches TIM-like sequences. This is visualized in Figure S4 by the increase of sequence motifs reported to be crucial for TIM activity (Olivares-Illana et al. 2017) throughout the generated sequences with progressive filtering. Further, sequences generated with the fine-tuned ProtGPT2 model do not follow the characteristic sequence length distribution of the fine-tuning dataset like ZymCTRL generated sequences do, but form a smooth distribution around the mean sequence length of the fine-tuning dataset (Figure S1B). Fine-tuned ProtGPT2 generated sequences show slightly higher sequence identities among each other compared to ZymCTRL (Figure S2). In contrast, the fine-tuned ProtGPT2 generated sequences are more distant to natural TIM proteins compared to ZymCTRL generated sequences but closer than sequences produced by the unconditional ProtGPT2 model (Figure S3).

To assess the ability of the fine-tuned ProtGTP2 to generate active TIMs, twelve promising variants, named gptTIMs, were selected for complementation assays to test TIM activity *in vivo*. All twelve proteins are classified as TIM by CATH, SCOPe, HHPred, and Foldseek and share a sequence identity of 33 % to 45 %. The gptTIMs are also diverse to the SpokenTIMs, sharing a sequence identity of mostly <30 % to the ZymCTRL generated TIMs (Figure S5). Two out of the twelve constructs showed rapid growth confirming that the novel proteins complement the desired TIM reaction (Figure 2B and Figure S6). Consequently, fine-tuning of unconditional models is a viable method to generate sequences performing a desired task.

### SpokenTIM9 is a well-folded enzyme with an efficiency close to natural TIMs

All SpokenTIMs and gptTIMs expressed well in *E. coli* BL21 at 37 °C albeit in varying amounts (Figure S7). We decided to experimentally characterize the best performing variant, SpokenTIM9, and to determine the biophysical properties of this LLM-designed enzyme. To avoid contamination from native *E. coli* TIM, SpokenTIM9 was purified to homogeneity in an *E. coli* BL21 Δtpi strain and assessed its biochemical properties. The far-UV circular dichroism spectrum of SpokenTIM9 confirmed the expected α/β-fold (Figure 3C). Intrinsic fluorescence of the aromatic residues revealed the putative TIM to possess a hydrophobic core (Figure 3D). SEC-MALS experiments at different protein concentrations showed a dynamic monomer-dimer equilibrium (Figure 3E). Individual peak analysis indicated monodisperse populations with a molecular weight in agreement to monomers (29.1 ± 0.4 kDa) and dimers (55.7 ± 0.9 kDa). However, the dimer peak indicated a dynamic population with a slight trend to higher oligomeric states. Different populations were visible in differential scanning calorimetry analysis. SpokenTIM9 unfolded irreversibly in two cooperative transitions with *T_m_*-values of 42.2 °C and 58.6 °C, respectively (Figure 3F). The presence of two transitions due to uncoupling of oligomer dissociation from the thermal unfolding of the monomer has been already observed for natural TIMs (Tellez et al. 2008), suggesting sequential transitions. In addition, the enthalpy change value indicates a folded protein with a well-packed core and a magnitude compatible with the expected value based on parametric equations (129.3 kcal mol^−1^ vs 182.2 kcal mol^−1^; (Robertson and Murphy 1997)).

**Figure 3.**
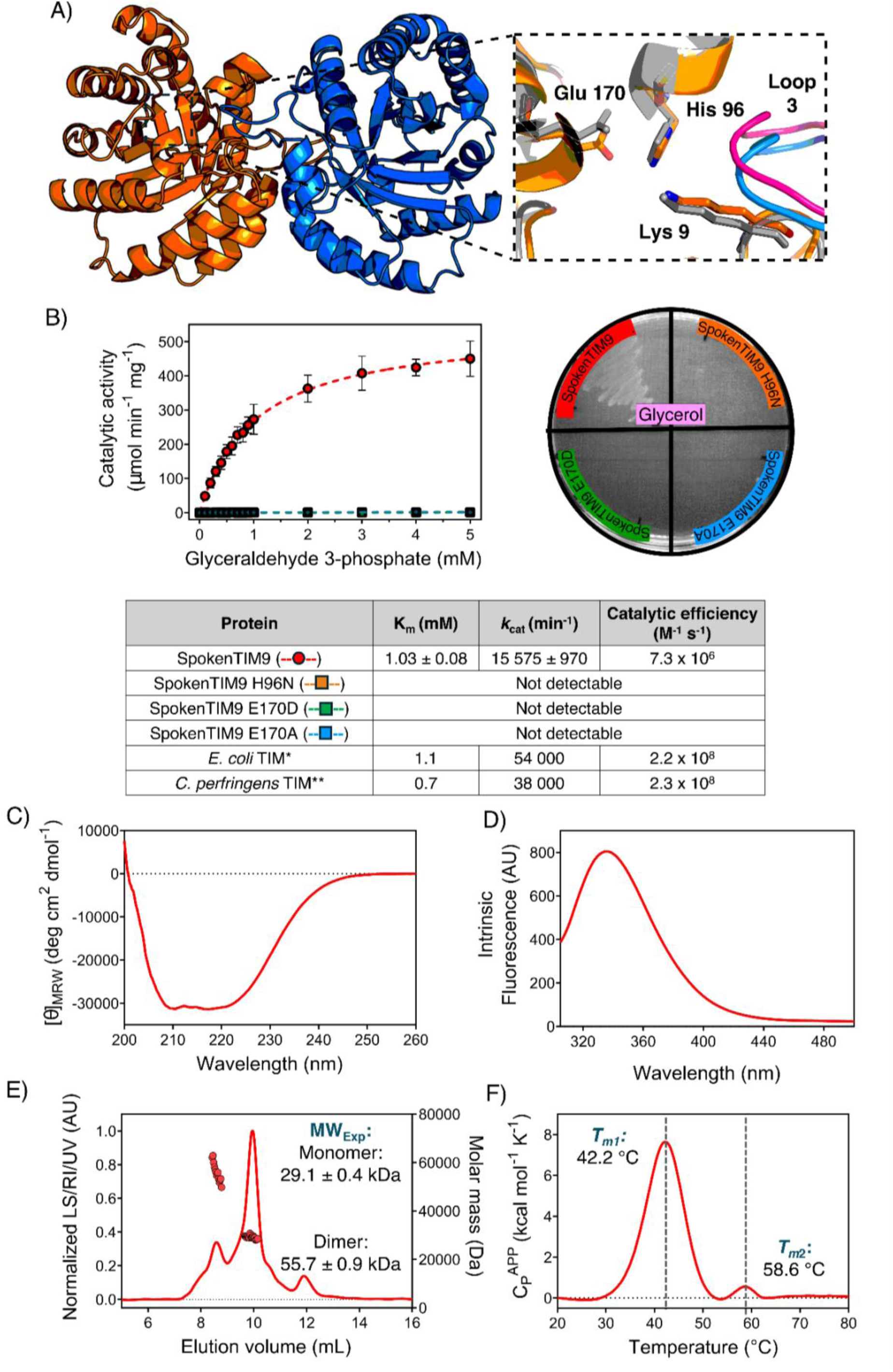
Biophysical and functional characterization of the artificial triosephosphate isomerase SpokenTIM9. (**A**) ColabFold model of the predicted dimeric structure of SpokenTIM9 with separate chains colored in orange and blue. Inset shows that the catalytic residues (orange) and the dimerization driving insertion loop (loop 3; blue) of SpokenTIM9 superpose very well with the corresponding residues of the bacterial TIM from *Clostridium perfringens* (catalytic residues: gray; loop 3: pink; PDB-ID: 4Y8F). (**B**) Catalytic activity of SpokenTIM9 was analyzed *in vitro* and *in vivo*. The activity of purified SpokenTIM9 is determined to be 7.3×10^6^ M^−1^s^−1^, which is 30-fold lower compared to *E.coli* TIM (*Alvarez et al. 1998) and *C.perfringens* TIM (**Romero-Romero et al. 2015) due to a lower *k*_cat_ value. Activity was further confirmed by complementation in *E. coli* Δtpi via growth on minimal media with glycerol as sole carbon source. Mutation of the active site residues H96 or E170 leads to loss of activity as well as the proteins’ ability to complement. (**C)** CD-spectrum of purified SpokenTIM9 confirms the expected ɑβ-fold. (**D**) Blue shift of the tryptophan fluorescence indicates SpokenTIM9 to be well-folded with a packed hydrophobic core. (**E**) SEC-MALS analysis revealed a monomer-dimer equilibrium, with the monomeric state being more populated. The mass of the monomer is determined to be 29.1 ± 0.4 kDa (polydispersity: 1.000) which is in agreement with the theoretical molecular weight of 28.9 kDa. The dimer shows a molecular mass of 55.7 ± 0.9 kDa (polydispersity: 1.004), matching with its theoretical molecular weight of 57.2 KDa. (**F**) DSC analysis of SpokenTIM9 shows the protein to unfold in a two-step process with T_m_-values of 42.2 °C and 58.6 °C, respectively.

Furthermore, enzyme activity was tested *in vitro* after purifying the protein from a Δtpi deletion strain. Experiments in biological triplicates using a coupled assay with glycerol-3-phosphate dehydrogenase (α-GDH) showed enzyme kinetics following a Michaelis-Menten behavior (red curve in Figure 3B) as has been observed for all natural TIMs characterized so far. The K_M_ value of SpokenTIM9 is in the same range for natural TIMs with the catalytic efficiency (*k*_cat_/K_M_) being just 30-fold lower than *E.coli* and *C.perfringens* TIM. This might suggest that its catalysis is not diffusion limited. However, SpokenTIM9 catalytic values are close to those reported for *Helicobacter pylori* as well as engineered triosephosphate isomerases (Table S4) such as ccTIM (consensus TIM; (Goyal et al. 2014; Sullivan et al. 2011)), MonoTIM (monomeric mutant; (Schliebs et al. 1996)), RMM0-1 (evolved from MonoTIM; (Saab-Rincón et al. 2001)), and LFYAA (stabilized interface by Rosetta; (Peimbert et al. 2008)). In contrast to the engineered variants, SpokenTIM9 is far off in sequence to natural TIMs and was designed in a single step.

To confirm the enzyme properties of SpokenTIM9, catalytic mutants (H96N, E170D, and E170A) were generated. These catalytic mutants showed no complementation in a TIM knock-out strain even after 7 days (Figure 3B and Figure S8). This was verified after purifying the protein variants and performing *in vitro* kinetic assays, thereby confirming the absence of catalytic activity. Overall, the results suggest that SpokenTIM9 TIM activity is indeed performed by the canonical catalytic residues as observed in natural TIMs. Additionally, catalytic mutants (H→N, E→D/A) of SpokenTIM4 and SpokenTIM5 were assayed *in vivo* on both carbon sources and also showed no growth in contrast to the original SpokenTIM constructs (Figure S8) supporting their activity being dependent on the same catalytic residues.

### LLMs recapitulate long-range interactions relevant for protein stability and function

Understanding the interactions within proteins is crucial for deciphering their structure, stability, and function. The proteins’ amino acids interact through various non-covalent and covalent forces, driving the protein to fold into its native three-dimensional structure, guiding catalysis, and dictating its interactions with other biomolecules. These interactions include hydrogen bonds, hydrophobic interactions, ionic bonds and van-der-Waals forces, which contribute to the fine-tuning of the overall shape of the protein (Pace et al. 2014). Therefore, it is remarkable that pLMs originally designed for the processing of human languages appear to also learn the *language of proteins* since they succeed in recapitulating such crucial interactions.

As illustrated with the predicted structure of our best performing protein, SpokenTIM9 (Figure 4), the pLM was not only able to recapitulate above-mentioned interactions but also place each residue precisely to create this active enzyme. In the predicted structure of SpokenTIM9 we highlight fundamental interactions like salt bridges, pi-stacking, hydrogen bonding networks and hydrophobic clusters such as the one in the core of SpokenTIM9 which is driven by eight residues throughout the entire sequence (Figure 4 and Figure S9). These interactions of residues throughout the sequence confer stability and participate drastically in shaping the desired TIM - barrel fold, which harbors a well-organized active site that promotes efficient catalysis *in vivo* and *in vitro*.

**Figure 4.**
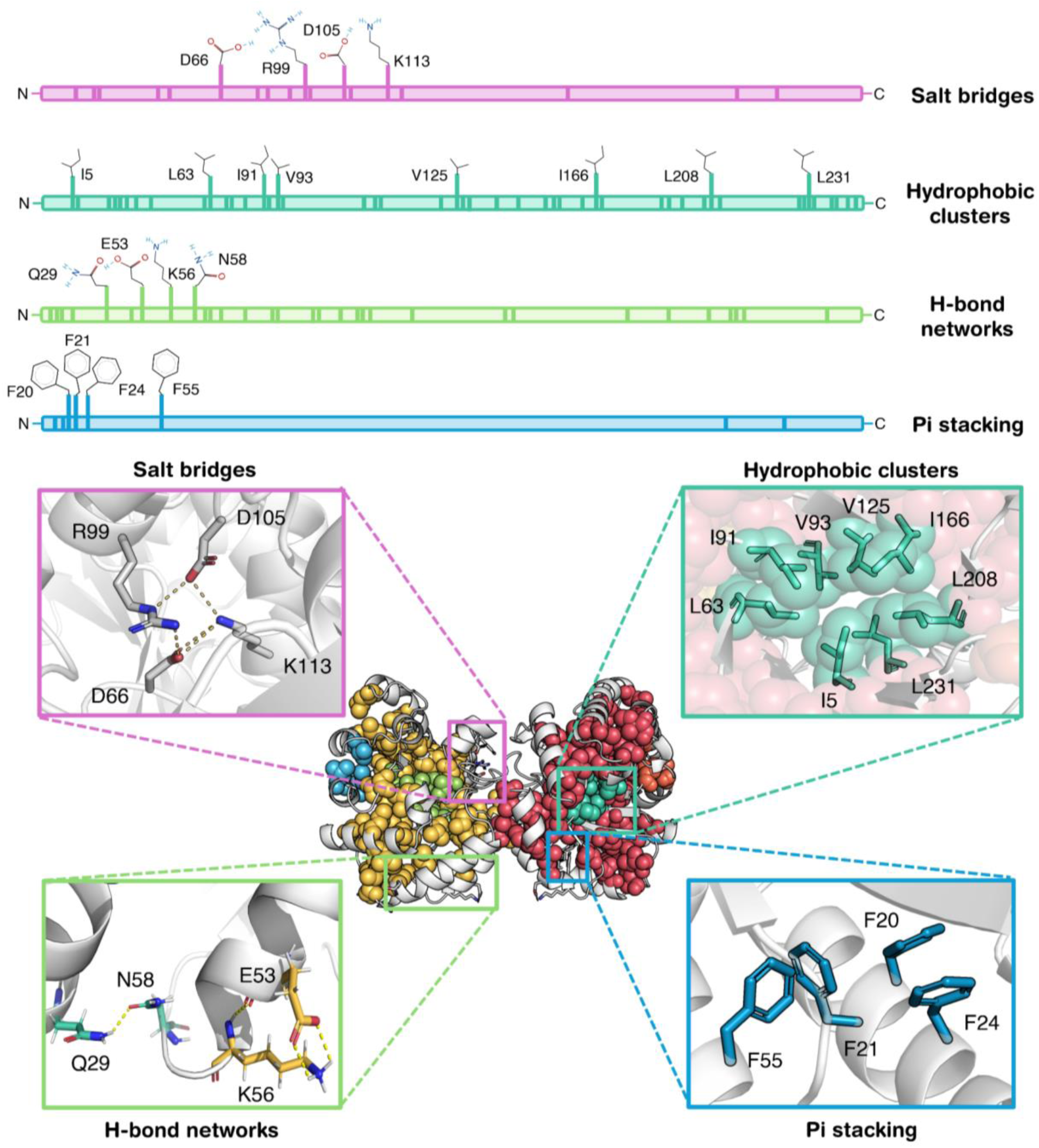
The predicted structure of SpokenTIM9 forms nature-like interactions throughout the entire sequence. The predicted model of the ZymCTRL generated and active SpokenTIM9 shows several amino acid interactions ranging throughout the entire length of the sequence, such as the formation of a hydrophobic core using ILV clusters (spheres, upper right), salt bridges (upper left), hydrogen-bonding networks (lower left), and aromatic interactions (lower right). Residues participating in the highlighted interactions are depicted as full side chains (top). Further residues participating in the respective kind of interaction throughout the sequence are marked as dashes at the proper position.

## Conclusions

There is a great need for tailor-made proteins and in particular the design of enzymes that carry defined properties. To reach this goal, we must gain a better understanding of the *language of proteins*. In recent years, the design of enzymes relied on physical methods to generate sequences folding into a given scaffold and implementing enzymatic activity via stabilization of a theozyme. Resulting activities were further enhanced by directed evolution. The introduction of AlphaFold2 in 2021, inspired novel AI-based methods as powerful tools for protein design. With tools like ProteinMPNN and RFdiffusion, the generation of novel scaffolds were tackled with unprecedented success. Nevertheless, the design of highly active complex enzymes is still a challenge to accomplish. Here we explored the capabilities of two pLMs, the conditional ZymCTRL and the unconditional ProtGTP2 that are based on different generation procedures, training datasets, and tokenization to design novel triosephosphate isomerases (EC 5.3.1.1).

ZymCTRL-generated sequences accurately followed the distribution of length and identities in the training dataset, indicating pLMs to be able to recapitulate a large diversity of properties inherent in the protein primary structure. Constructs for experimental characterization were built by conditional generation followed by shallow filtering based on sequence length, a low sequence identity to any natural protein (≤ 35 %), and visual inspection of ColabFold predictions concerning the presence and orientation of catalytic residues and the predicted oligomeric state. Three out of twelve designs were found to be active *in vivo* with SpokenTIM9 being the most promising variant. SpokenTIM4 and SpokenTIM5 grew more slowly in the complementation assay likely due to lower activity or solubility. The activity of all three active SpokenTIMs was confirmed by assaying constructs with mutated catalytic histidine or glutamate, which showed no growth even after 7 days of incubation.

Next, we investigated the ability of the unconditional language model ProtGPT2 to generate active enzymes after fine-tuning the model. The ProtGPT2-generated sequences were filtered stringently. Successful filtering showed a rapid increase of desired sequence motifs. Unlike ZymCTRL-generated sequences, fine-tuned ProtGPT2-generated sequences do not follow trends like sequence length and diversity of the training/fine-tuning dataset. Of the twelve gptTIMs generated with the fine-tuned model and chosen for experimental characterization, two variants showed *in vivo* complementation in a TIM-deficient strain. This shows that both pLMs, even though they differ in generation, training, and tokenization, are able to design proteins with the desired enzyme activity.

Biochemical characterization of the best performing construct, SpokenTIM9, revealed that the artificial protein is well folded and possesses biophysical parameters similar to natural TIMs. However, SpokenTIM9 adopts a dynamic monomer/dimer equilibrium and shows *in vitro* activity around 30-fold lower than natural TIMs. It was confirmed that activity is based on the canonical catalytic residues as their mutation leads to complete loss of activity. The behavior of SpokenTIM9 indicates that the generated protein is not yet perfect. However, it is remarkable that the pLMs were able to generate such complex and efficient enzymes that are active *in vivo*. Moreover, the tested sequences are far more distant to natural counterparts than shown for other pLM generated sequences so far, illustrating the potential for diversification and further evolution or engineering to include new features in those proteins in the future.

Taken together, we showed the application of two different LLMs to generate complex active enzymes, demonstrating pLMs to be powerful tools for protein design. These novel tools provide an innovative way to explore sequence space and the opportunity to gain deeper insights into the *language of proteins*.

## Materials and Methods

### Sequence generation, analysis, and filtering

92,379 sequences were generated using ZymCTRL (Munsamy et al. 2024) with the enzyme commission number EC 5.3.1.1 as conditional input prompt. Generation was conducted with 20 sequences per batch with a maximum sequence length of 1024 amino acids. Repetition penalty was set to 1.2, Top_k was set to 8, and the temperature parameter was set to 1.

Fine-tuning of the ProtGPT2 model (Ferruz et al. 2022) was performed over one epoch with a learning rate of 0.0001 without warm-up ramping using 940 randomly selected natural triosephosphate isomerase sequences, including all previously experimentally characterized ones. Then, 14,300 sequences were generated by 100 sequences per batch without length restriction, repetition penalty value set as 1.2 and Top_k 9.

Sequence identity to natural proteins was determined using BLASTp from the BLAST+ v2.10.1 software (Camacho et al. 2009). BLAST searches were performed against the NCBI RefSeq90 sequence database. Sequence comparison among generated sequences was carried out by performing an all-vs-all alignment of the respective sequence datasets using MMseqs2 (Mirdita et al. 2019; Steinegger and Söding 2017) with a minimum coverage of 60% (−*c 0.6, --cov-mode 0*) and an E-value cutoff of 0.001 (*-e 0.001*). Sequence alignments were performed using MAFFT (Nakamura et al. 2018) with a gap opening penalty of 1.53 (*--op 1.53*) and a gap extension penalty of 0.123 (−-ep 0.123) using the FFT-large-NS-2 alignment strategy (automatically selected via *--auto*). HMM profiles were created using the *hmmbuild* command of the HMMER3 suite package (Eddy 2011). Consensus columns were defined (*--fast*) to be populated with more or equal to 50 % residues opposed to gaps (*--symfrac 0.5*). Letter heights were given by the hmmlogo command discarding glyphs below background (*--height_relent_abovebg*). Logos were illustrated using the *logomaker* python module. CATH classification of selected generated proteins was assigned by using the respective sequences as input for the CATHdb sequence search tool (Sillitoe et al. 2021). SCOPe classifications (Fox et al. 2014) were determined by using the respective sequences as input for the Fuzzle2.0 web server (Ferruz et al. 2021a).

Structures were modeled using AlphaFold2 ColabFold Batch v.1.3.0 (Mirdita et al. 2022) without amber relaxation. Sequences without His_6_-tag were used as input. Oligomeric states were modeled using ColabFold Batch v1.3.0 using fasta-files containing the sequence of choice in the multiplicity of the desired oligomeric state with the monomeric sequences being separated by a “:” character. ColabFold models of the respective proteins were used as input for a structure-based protein comparison with Foldseek (Van Kempen et al. 2023). Hydrophobic solvent-accessible surface area of the ColabFold models was determined with the FreeSASA python library. The ideal surface area of the proteins was calculated according to (Dill et al. 2011) with the ideal radius 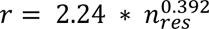, where *n_res_* is the number of residues. The spatial aggregation propensity of the ColabFold models was analyzed using the *SapScoreMetric* function from RosettaScripts (Fleishman et al. 2011) from the Rosetta Commons Software Suite v3.13 (Leaver-Fay et al. 2011). Mean charge of a given sequence was calculated using the *SequenceParameters* function of the localcider python library. ILV clusters were determined and depicted using the ProteinTools web server (Ferruz et al. 2021b).

### Reagents and constructs

Analytical-grade chemicals were used for all experiments. Water was distilled and deionized. All SpokenTIM and gptTIM sequences were obtained from BioCat and directly cloned into pET21b(+) using the 5’ NdeI and 3’ XhoI restriction sites. Catalytic mutants for SpokenTIMs were generated via QuikChange-PCR (Agilent) with templates from wild-type genes and proper mutation primers, and using *E. coli* Top10 cells (Invitrogen) for cloning. Sequencing was performed to confirm the individual point mutations.

### Protein overexpression and purification

LB-media pre-cultures supplemented with the antibiotic resistance were inoculated with a single colony of the respective construct in *E. coli* BL21(DE3) or *E. coli* BL21(DE3) Δtpi. 1 L TB-media expression cultures were inoculated and grown at 37 °C until an OD_600_ of 0.6 – 1.0 was reached, whereupon overexpression was induced with 1 mM IPTG. Cells were harvested after 18-20 hours of expression, washed with ddH_2_O and frozen at −20 °C until further use.

For SpokenTIM9, pellets were resuspended in 20 mM TEA pH 8.0, 300 mM NaCl, 20 mM Imidazole, and 1 mM TCEP buffer, with the addition of SERVA protease inhibitor mix. Lysis was conducted by addition of 10 mg hen egg lysozyme (Sigma Aldrich) incubated at 37 °C for 20 minutes followed by sonication. Lysed cells were centrifuged at 40,000 g for 1 hour. The filtered supernatant was applied to a Ni^2+^-IMAC column (HisTrapHP 5 mL, Cytiva), washed with 20 mL lysis buffer and eluted using an gradient from 0 % - 60 % of high imidazole buffer (20 mM TEA pH 8.0, 300 mM NaCl, 500 mM imidazole, 1 mM TCEP). Fractions containing the protein of interest were dialyzed exhaustively against 10 mM TEA pH 8.0, 1 mM EDTA, 1 mM TCEP and purified by anion exchange chromatography (MonoQ 5/50 GL 1 mL, Cytiva). Elution was performed with a gradient from 0 % to 60 % of high salt buffer (10 mM TEA pH 8.0, 1 mM EDTA, 1 M NaCl, and 1 mM TCEP). Fractions containing the SpokenTIM construct were applied to a SEC column (HiLoad Superdex 75 pg, Cytiva) using 10 mM TEA pH 8.0, 1 mM EDTA, 1 mM TCEP, and 150 mM NaCl as running buffer. Fractions corresponding to the estimated dimeric species were pooled and used for further experiments.

Protein identity was confirmed by mass spectroscopy. Protein samples were analyzed by SDS-PAGE. Bands corresponding to the protein of interest were excised from the gel and trypsinized according to (Shevchenko et al. 2010). SpokenTIM9 was identified via LC ESI-MS/MS on an LTQ-XL mass spectrometer coupled to an EASY-nLC II chromatographic system according to (Weiss et al. 2022).

### Bacterial complementation in an E. coli Δtpi strain (in vivo activity)

*E. coli* BL21(DE3) Δtpi cells were transformed with the respective constructs and grown in LB-media. Cells were pelleted at an OD_600_ mL^−1^ ratio of 3 and washed twice with 1x PBS buffer. The pellet was resuspended in 100 µL 1xPBS and 20 µL were plated on M9 minimal media agar plates supplemented with 100 µg mL^−1^ uracil, 1 µg mL^−1^ thiamin, 40 µg mL^−1^ histidine, and 0.4 % of either glucose (w/v) or glycerol (v/v) as the sole carbon source. Plates were incubated at 37 °C and photo-documented every 24 hours. As a positive control the bacterial TIM from *Clostridium perfringens* (Romero-Romero et al. 2015), the eukaryotic TIM from *Schizosaccharomyces pombe* (Romero-Romero and Garza-Ramos 2021), and the archaeal TIM from *Methanotorris formicicus* were used as positive controls. An empty pET-28b(+) vector was used as a negative control. Plates were inspected every 24 hours for six days. During this time the negative control did not show any growth of bacterial colonies.

### In vitro enzymatic assay

TIM activity from GAP to DHAP was determined using a coupled assay with glycerol-3-phosphate dehydrogenase (α-GDH) (Romero-Romero et al. 2015). Enzymatic activity was measured in 100 mM TEA, 10 mM EDTA, 1 mM DTT pH 8.0 supplemented with 20 µg α-GDH and 0.2 mM NADH. The catalytic assay was measured at 25 °C and 250 ng mL^−1^ TIM concentration by varying substrate (GAP) from 0.1-5.0 mM. Activity was monitored by the reduction in absorption at 340 nm in a Genesys 10S UV-Vis Spectrophotometer (Thermo Scientific) and considering only the linear velocities. Raw data were converted to specific activity using the extinction of NADH (ε_340_= 6220 M^−1^ cm^−1^), a path length of 1 cm, and the enzyme concentration in the assay:

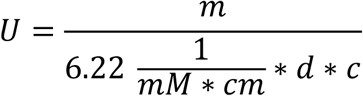

where *U* represents the specific activity at a given GAP concentration in 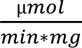, *m* the slope of the linear decrease resulting from NADH oxidation in 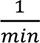, *d* the path length, and *c* the final protein concentration in 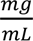. Experimental data were fitted using the Michaelis–Menten equation (Cornish-Bowden 2012):

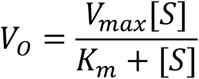

where *V*_0_ represents the initial velocity, [S] the substrate concentration, *V*_max_ the maximum velocity in the assay, and K_m_ the concentration of substrate at which the reaction rate is half of *V*_max_. The catalytic efficiency (*k*_cat_/K_m_) was calculated by adjusting the K_m_ value considering that only 4 % of GAP in aqueous solution is in the unhydrated aldehyde substrate, the only form that TIM catalyzes (Trentham et al. 1969).

### Biophysical characterization of SpokenTIM9

For far-UV circular dichroism (CD) measurements proteins were dialyzed exhaustively against 10 mM sodium phosphate pH 8.0 and 100 mM NaCl buffer. Measurements were performed at 25 °C using a protein concentration of 0.2 mg mL^−1^ (J-710 Spectropolarimeter). CD data were converted to mean residue ellipticity:

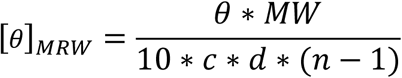

where [*θ*]*_MRW_* represents the mean residue ellipticity in 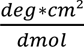, *θ* the ellipticity in mdeg, *MW* the molecular weight in 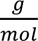, *c* the protein concentration in 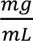, *d* the cuvette path length in cm, and *n* the number of amino acids in the protein.

For intrinsic fluorescence (IF) proteins were dialyzed exhaustively against 10 mM sodium phosphate pH 8.0 and 100 mM NaCl buffer. IF spectra were collected with excitation wavelengths of 280 nm and 295 nm, the latter solely exciting tryptophan residues. Measurements were performed at 25 °C using a protein concentration of 0.2 mg mL^−1^ (FP-6500 Spectrofluorometer). Fluorescence spectra were normalized considering the intensity maximum for each experiment. Fluorescence spectral center of mass (SCM) was calculated from intensity data at each scan (Iλ) obtained at different wavelengths (λ): SCM= ∑λIλ/∑Iλ.

For Size Exclusion Chromatography coupled Multi Angle Light Scattering (SEC-MALS), protein samples were applied to a Superdex S75 Increase 10/300 column coupled to a 1260 Infinity II HPLC system and a miniDAWN MALS detector with an Optilab differential refraction index detector. Samples were run with a flow rate of 0.8 mL min^−1^ using 10 mM TEA pH 8.0, 1 mM EDTA, 1 mM TCEP and 150 mM NaCl buffer as mobile phase. For normalization, peak alignment, and band broadening, a bovine serum albumin standard with a concentration of 2 mg mL^−1^ was run before and after a set of measurements (Pierce™ Bovine Serum Albumin Standard Ampules 2 mg mL^−1^). Data were processed and evaluated using the ASTRA 8.0.2.5 software.

For Differential Scanning Calorimetry (DSC) experiments, proteins at 1, 2 and 3 mg mL^−1^ were exhaustively dialyzed against 10 mM sodium phosphate pH 8.0, 100 mM NaCl buffer then degassed at room temperature. Measurements were performed in a MicroCal Peaq-DSC system (MicroCal®, Malvern Panalytical) with a scanning speed of 90 K h^−1^ in a non-feedback mode. In order to have proper instrument equilibration, two buffer–buffer scans were performed before each protein-buffer scan. The corresponding buffer–buffer trace was subtracted from each endotherm and also the chemical baseline for each scan, which was then normalized considering the protein concentration in each experiment. MicroCal software was used for data analysis.

## Acknowledgments

We thank Gloria Saab Rincón for the donation of the *E. coli* BL21 Δtpi strain, acknowledge Sooruban Shanmugaratnam, Susanne Schäfer and Sabrina Wischt for competent technical support. Support from the Elite Network of Bavaria and its study program “Biological Physics” is gratefully acknowledged by AEB and BH.

## Funding

This work was supported through core funding of the University of Bayreuth. We further acknowledge financial support from the European Union’s Horizon 2020 research and innovation programme under grant agreement No 951375 (ArtMotor).

## Competing interests

The authors declare that they have no conflicts of interest with the contents of this article.

## Author Contributions

SRR and BH designed research. NF generated ZymCTRL sequences. SRR and AEB generated ProtGPT2 sequences. SRR, AEB and SS analyzed sequences. AEB generated structural models. SRR, AEB and BH selected constructs for experimental testing. SRR, AEB and TK performed complementation assays. SRR and AEB expressed and purified proteins, performed enzyme kinetics and biophysical characterization. SRR, AEB and BH analyzed the data. SRR, AEB and BH wrote the manuscript. All authors discussed and commented on the manuscript.

## Data and materials availability

All data to support the conclusions of this manuscript are included in the main text and supplementary materials.

## Supplementary Materials

This article contains supplementary material that includes: supplementary figures 1-9 and supplementary tables 1-4.

## Abbreviations

CD: Circular Dichroism
DSC: Differential Scanning Calorimetry
IF: Intrinsic Fluorescence
MALS: Multi Angle Light Scattering
SEC: size exclusion chromatography
*T_m_*: midpoint of thermal unfolding
K_M_: Michaelis-Menten constant
*k*_cat_: catalytic constant
TIM: triosephosphate isomerase.

## Supplementary Information

**Figure S1.**
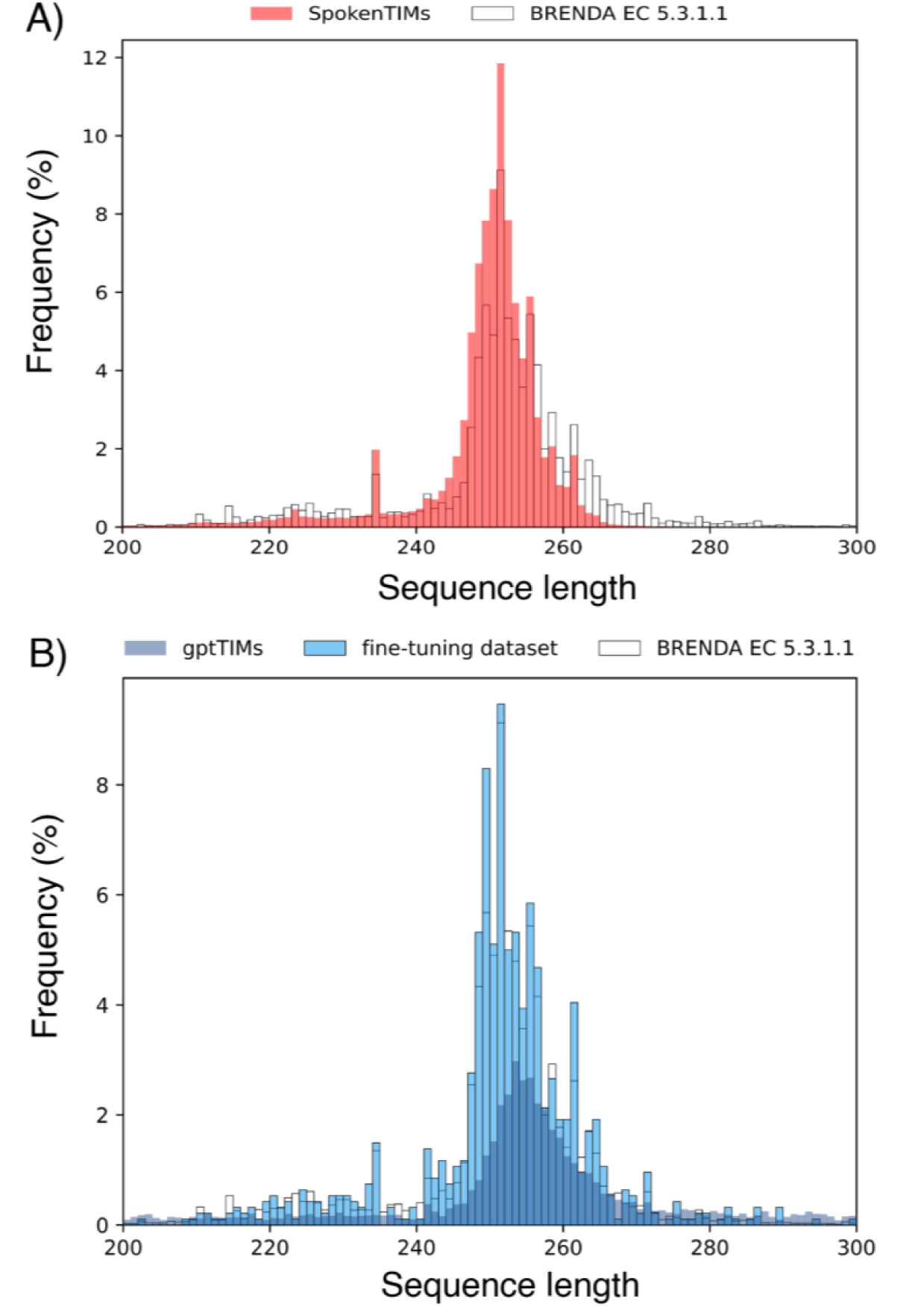
Protein language models recreate characteristic sequence lengths of the training datasets. **A)** The distribution of sequence lengths of TIM sequences generated with ZymCTRL using EC 5.3.1.1 as conditional input term (red) are superposed with the sequence length distribution of all sequences classified as triosephosphate isomerases (white). The sequences generated by ZymCTRL mimic the distribution of the sequences found in the training dataset with a major population at approximately 250 amino acids in length. **B)** ProtGPT2 fine-tuned on 940 natural triosephosphate isomerase sequences generates putative enzymes with a sequence length distribution (blue) centered around the mean sequence length of TIMs in the fine-tuning dataset (light-blue, 257 amino acids). The fine-tuning dataset mimics the characteristic length distribution of the sequences classified as TIMs (white).

**Figure S2.**
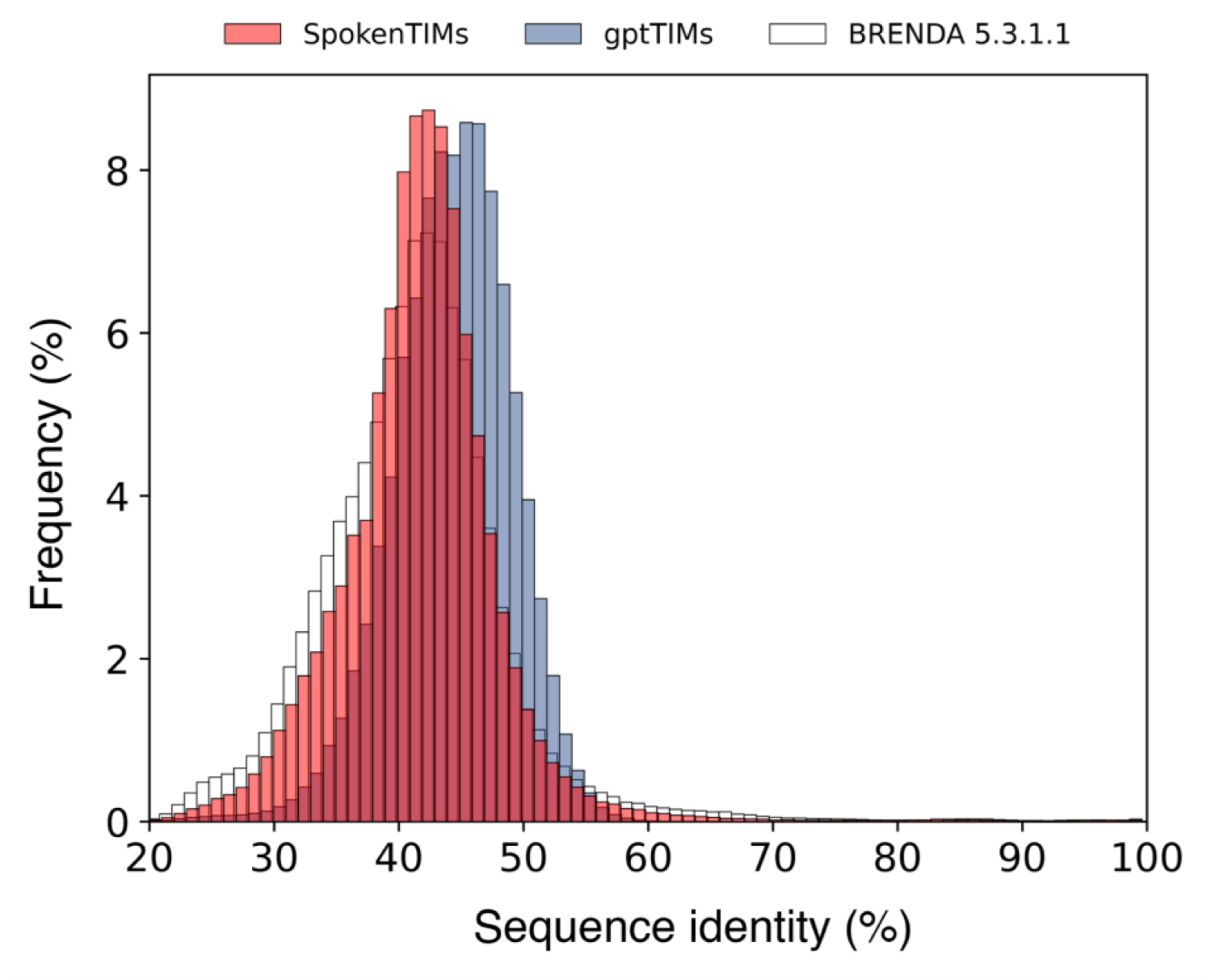
ZymCTRL and ProtGPT2 generated TIM sequences are highly diverse. Triosephosphate isomerase sequences generated with ZymCTRL (red) or fine-tuned ProtGTP2 (blue) are highly diverse, possessing sequence identities to the other generated sequences mostly lower than 50 %. The sequence identity distribution among ZymCTRL generated sequences follow closely the sequence identities of proteins classified as TIMs (white). The distribution of ProtGPT2 generated sequences is shifted slightly to higher identities. Sequence identities were created with an all-vs-all local alignment using MMseqs2.

**Figure S3.**
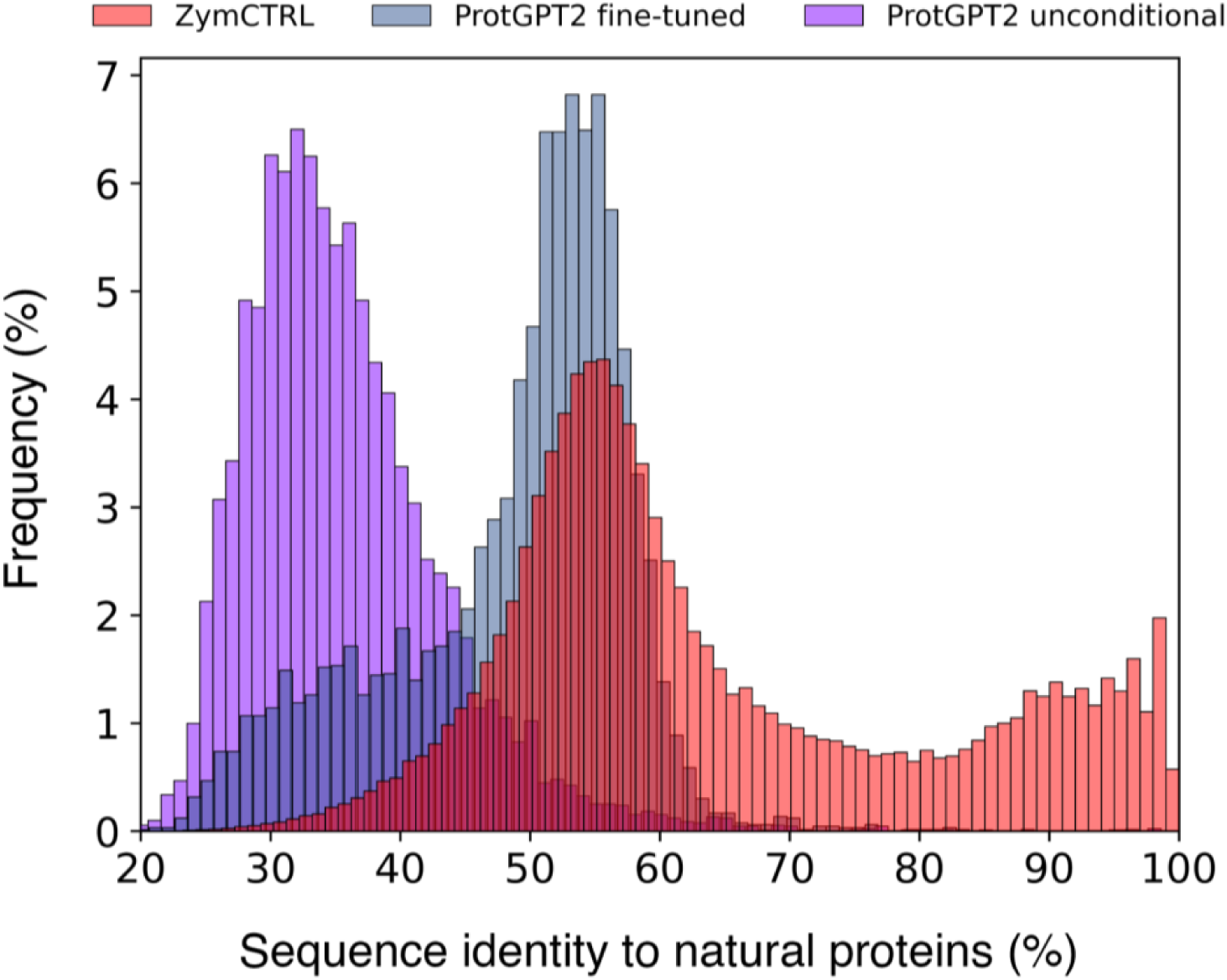
Language models generate sequences distinct from natural proteins. Triosephosphate isomerase sequences generated with ZymCTRL (red), the TIM-fine-tuned ProtGPT2 (blue), and *de novo* protein sequences generated with the unconditional ProtGPT2 model (purple) share mostly low sequence identity with natural proteins. ZymCTRL generated TIM sequences display most sequence identity to natural proteins with major populations at approx. 55 % and 90 % sequence identity. ProtGPT2 generated TIMs are more distinct with almost all sequences sharing less than 60 % identity to natural proteins. Nonetheless, random protein sequences generated with the non-fine-tuned ProtGPT2 model are most distinct from natural proteins, highlighting the influence of the fine-tuning on the generation process. Sequence identities to the best-hit natural protein were determined using BLASTp.

**Figure S4.**
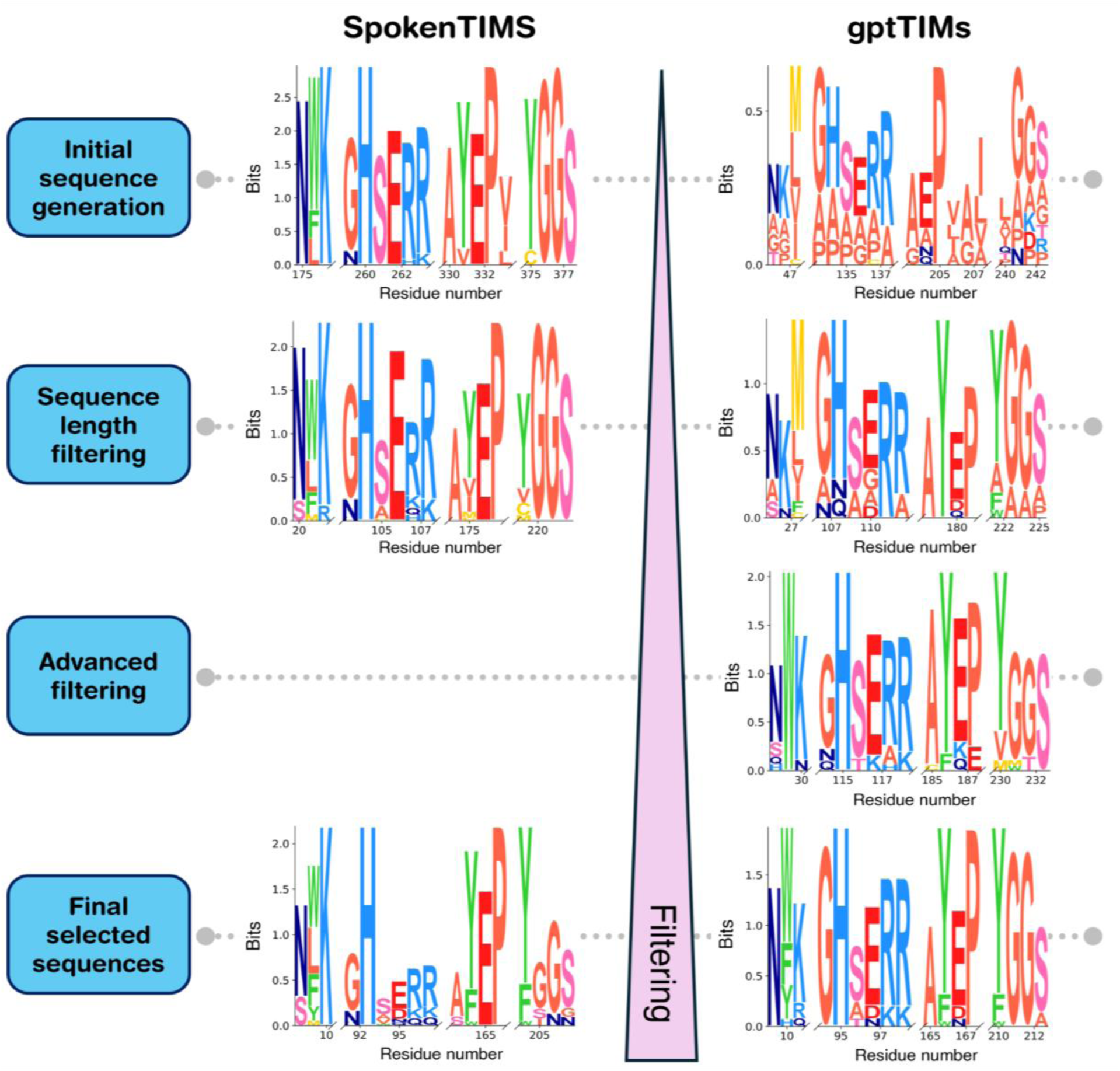
Filtering of LLM generated sequences improves average sequence performance. Sequence motifs were generated from profile hidden markov models based on multiple sequence alignments of all sequences present in the respective processing step. The motifs corresponding to the three catalytic residues (NW**<u>K</u>**, G**<u>H</u>**SERR, AY**<u>E</u>**P) and the loop closing motif (YGGS) are shown. ZymCTRL generated sequences show good depiction of the motifs without additional filtering. ProtGPT2 generated sequences are further away from coinciding on the motifs necessary to perform the desired TIM reaction. After mere sequence length filtering the representation of the necessary motifs increases. Sequences after the advanced filtering possess a sequence identity to any natural protein of ≤50 %. Nonetheless, sequence independent advanced filtering, based solely on surface hydrophobicity parameters and the average PAE value of the predicted models, results in large improvements of the average sequence performance as the desired motifs are clearly emerging.

**Figure S5.**
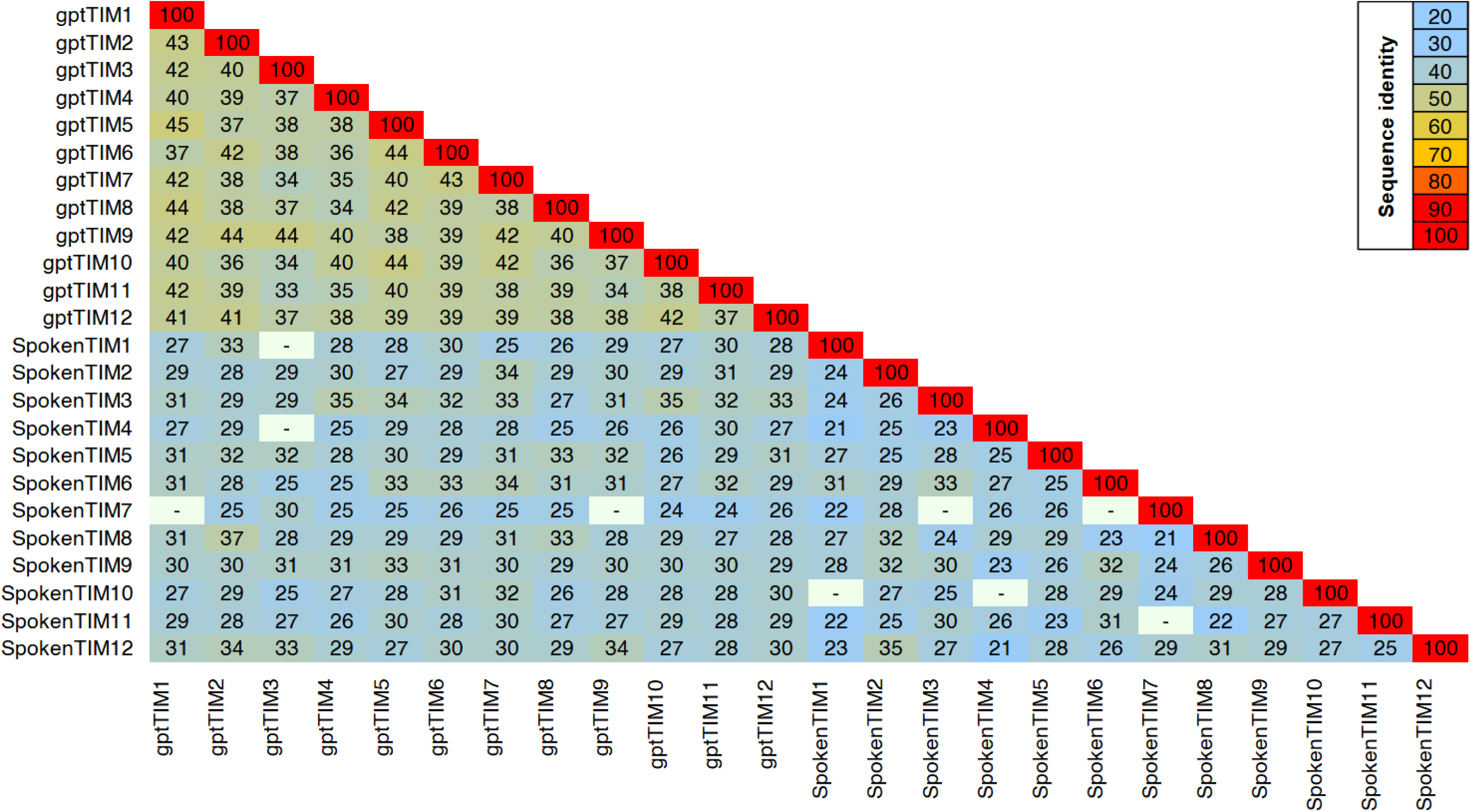
Sequence identity matrix between SpokenTIMs and gptTIMs. Sequence comparison was performed using MMseqs2. “-”: *e-value* of alignment > 0.001; numbers indicate identity in %.

**Figure S6.**
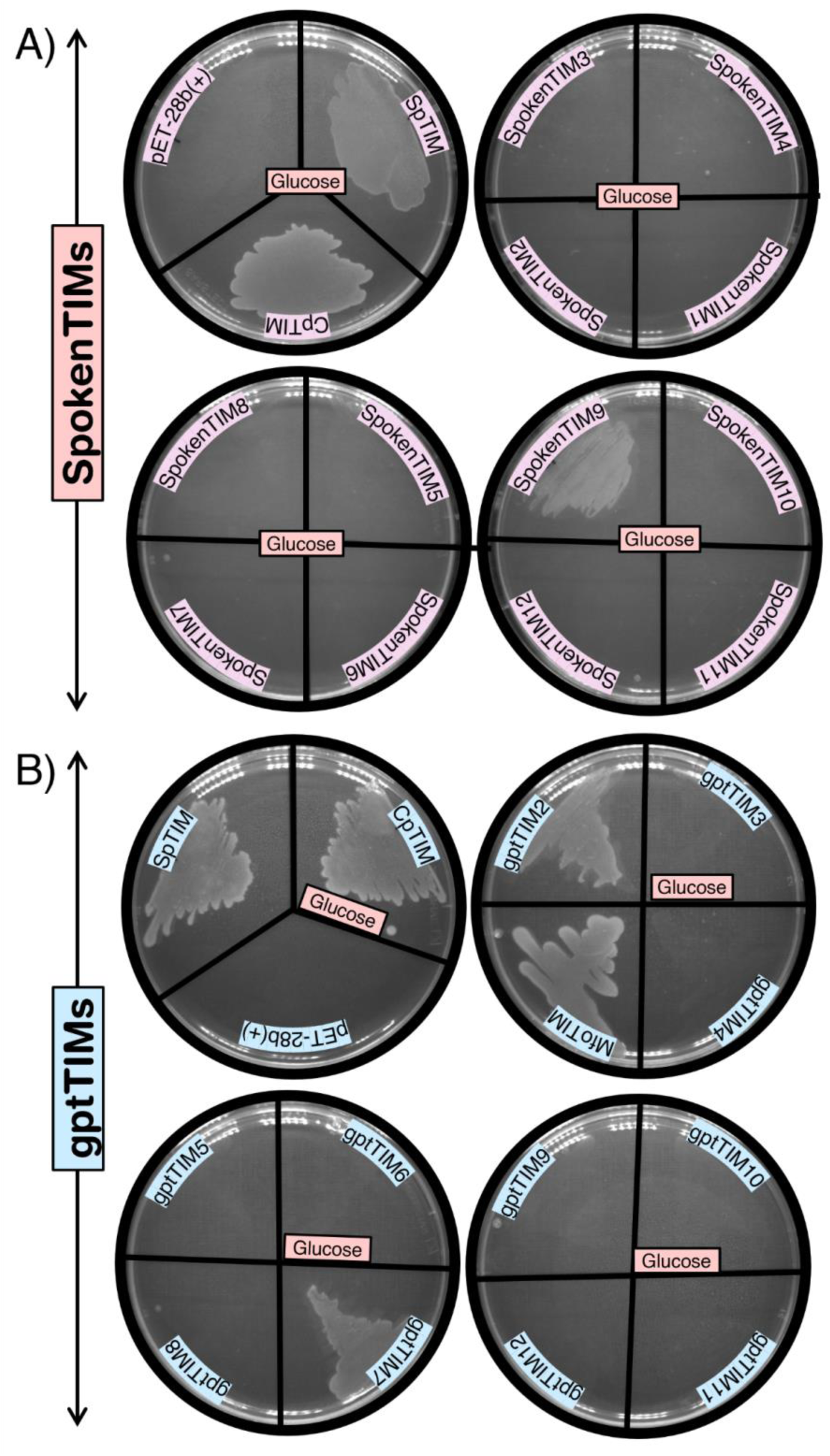
Bacterial complementation assay of SpokenTIMs and gptTIMs using glucose as carbon source. *E. coli* BL21 cells lacking the native gene for triosephosphate isomerase, but carrying the SpokenTIM or gptTIM constructs, were plated on M9 minimal media containing glucose as the sole carbon source. (**A**) Snapshot after 48 hours: SpokenTIM9 activity complements the gene-deficiency resulting in increased growth rates in comparison to cells carrying inactive/insoluble proteins or an empty pET28b(+) vector. Also SpokenTIM4 and SpokenTIM5 show slight growth. (**B**) gptTIM2 and gptTIM7 also show increased growth rates on glucose after 48 hours compared to cells carrying inactive proteins or an empty pET28b(+) vector. As positive controls cells carrying the natural TIMs from *Clostridium perfringens, Schizosaccharomyces pombe,* and *Methanotorris formicicus* were used.

**Figure S7.**
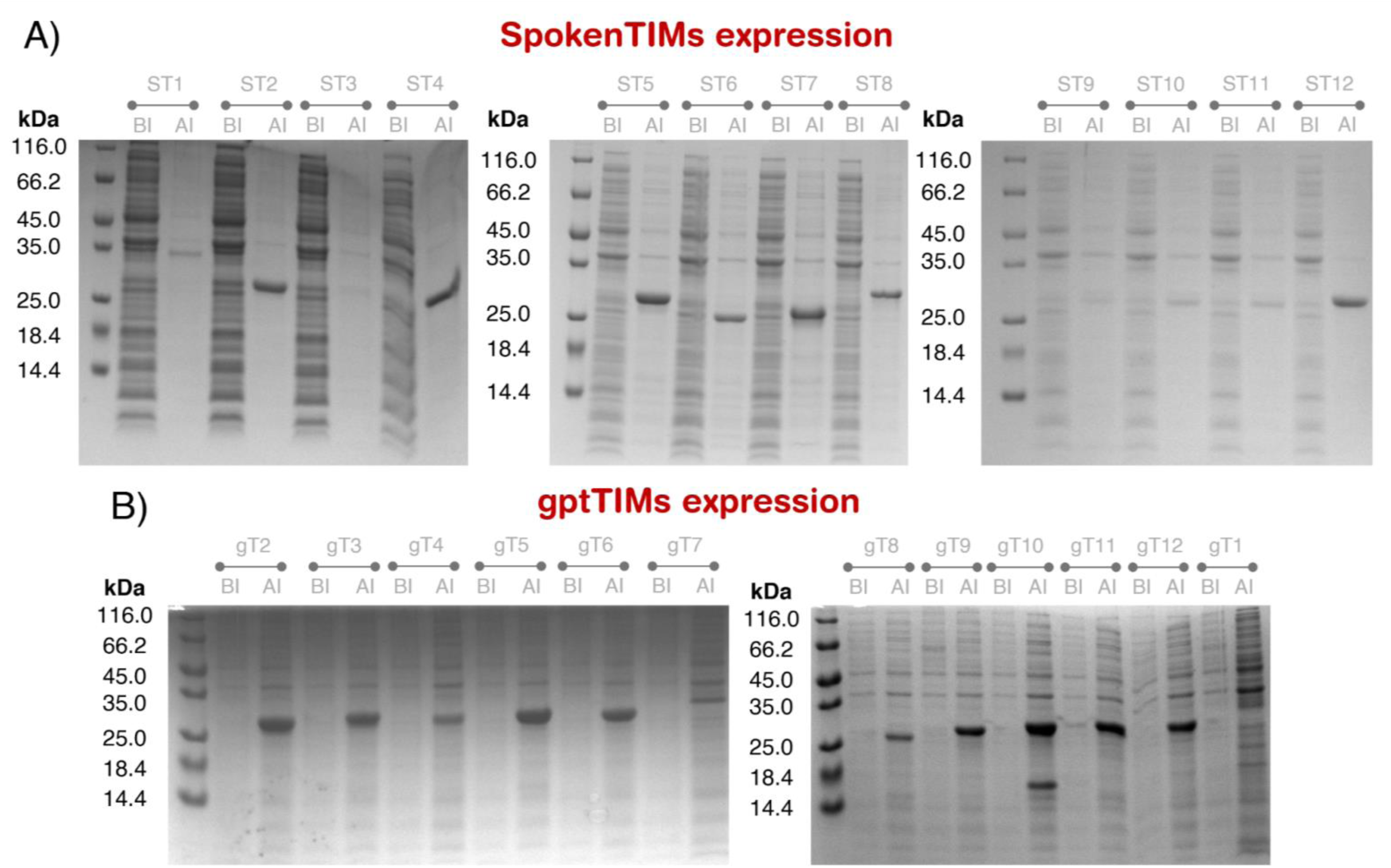
Expression gels of SpokenTIMs (A) and gptTIMs (B). Before (BI) and after induction (AI) SDS samples of SpokenTIM (STX; X = number of construct) and gptTIM (gTX) expressions. Expression was conducted in *E. coli* BL21(DE3) cells carrying the respective plasmid and induced with a final concentration of 1 mM IPTG. All artificial TIMs express in varying amounts and run at the expected size of approx 28 kDa.

**Figure S8.**
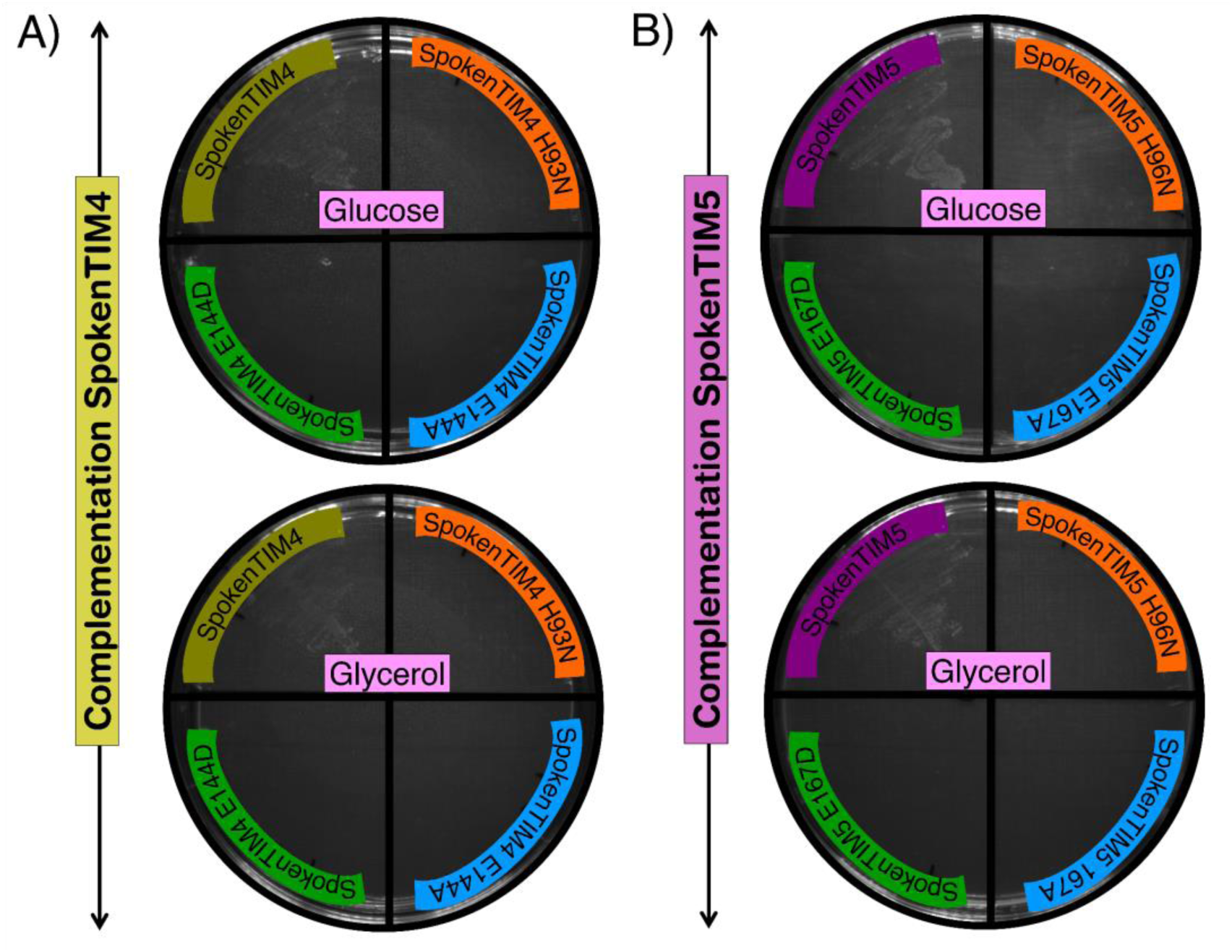
Bacterial complementation assay of the catalytic mutants of SpokenTIM4 and SpokenTIM5 using glucose or glycerol as carbon source. Activity was further confirmed by complementation in *E. coli* Δtpi via growth on minimal media with glucose or glycerol as sole carbon source. Mutation of the active site residues H or E leads to loss of activity as well as the proteins’ ability to complement.

**Figure S9.**
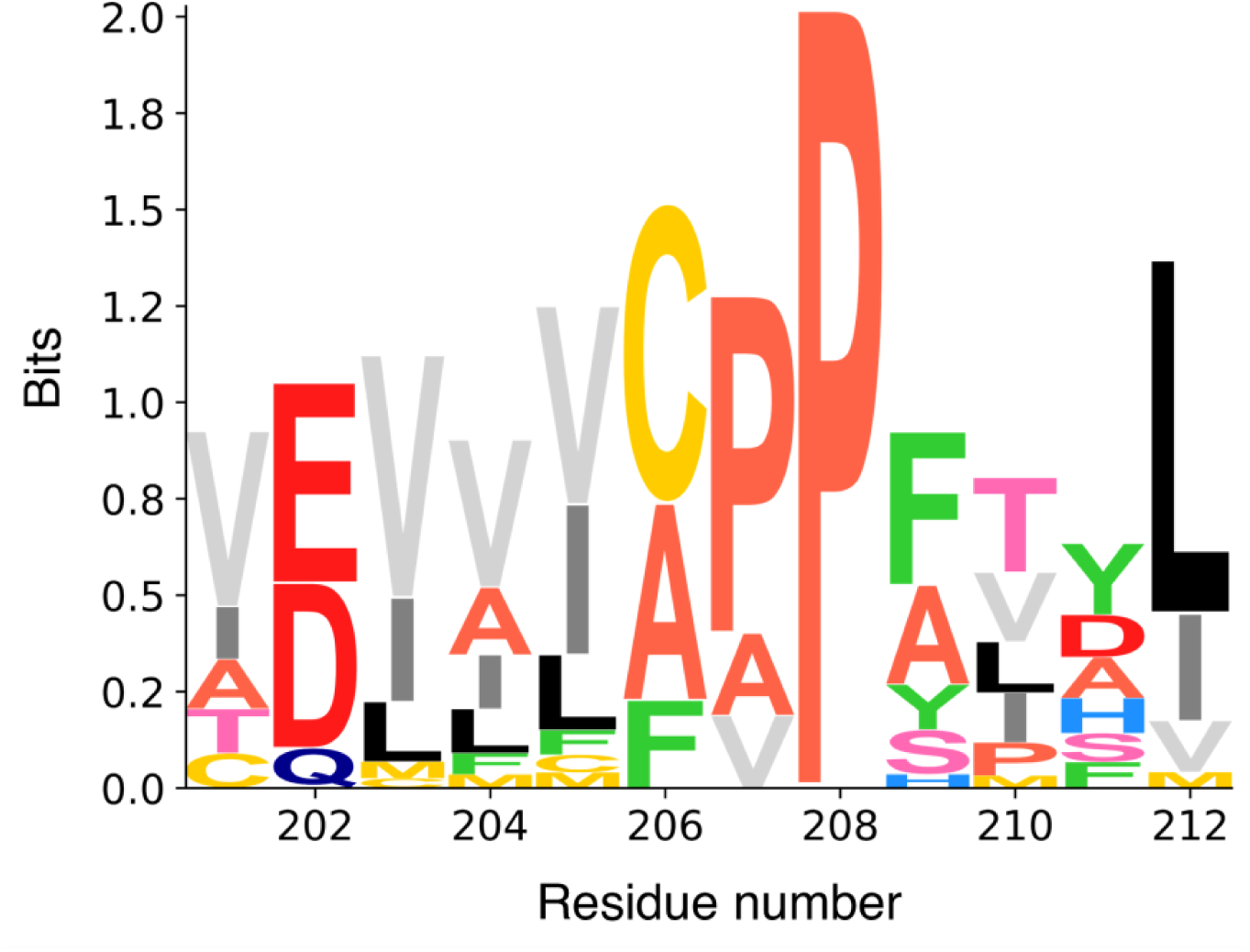
Protein LLMs recapitulate hydrophobic clusters. The sequence motif was generated from an HMM based on a multiple sequence alignment of all generated ZymCTRL sequences. Valine (light gray), leucine (black), and isoleucine (dark gray) are used almost interchangeably in the generation, indicating the language model recapitulating the interactions needed for the formation of the hydrophobic core and stability mitigating ILV cluster found throughout the predicted protein structures of the generated putative TIMs.

**Table S1.**
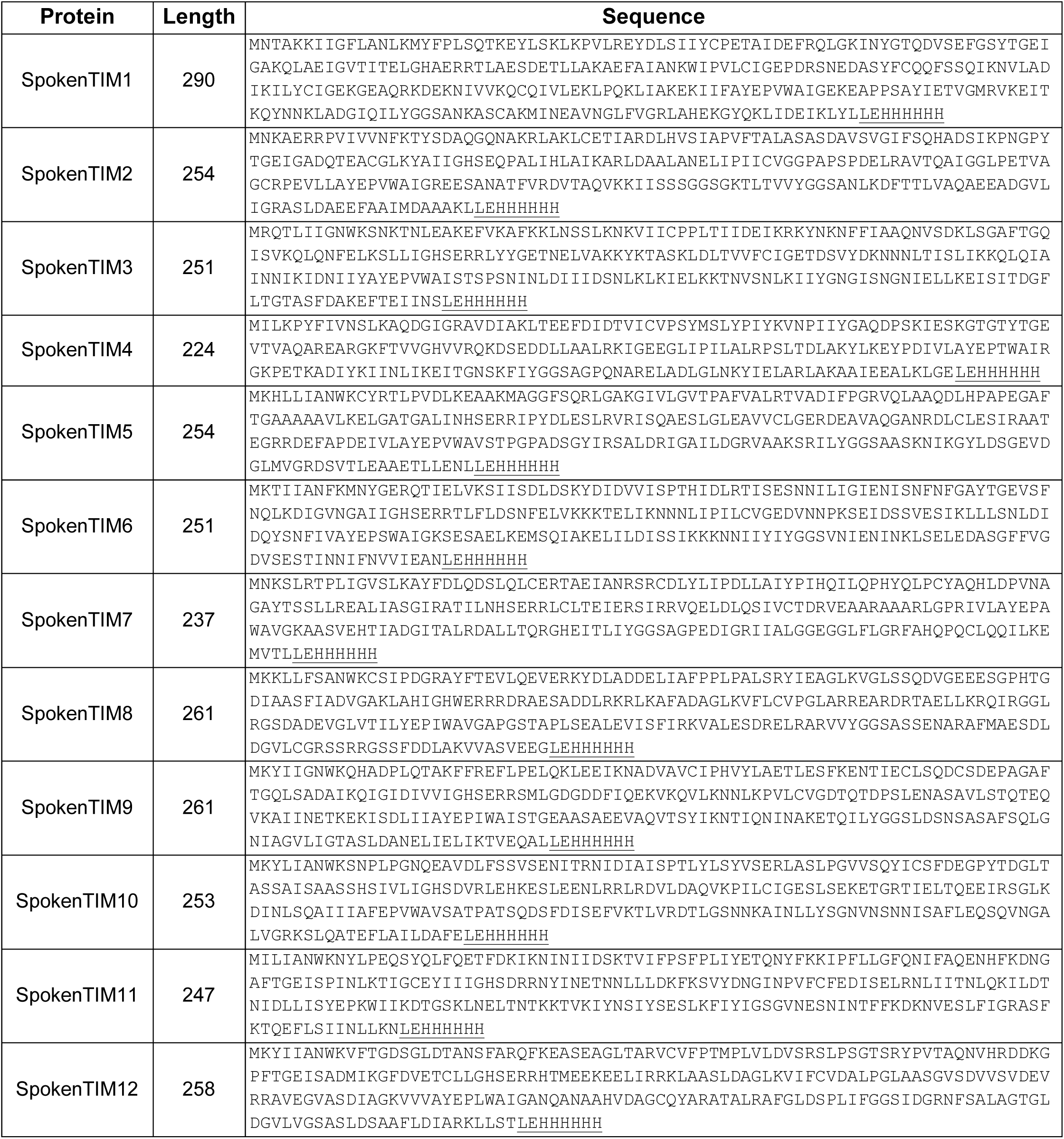
Amino acid sequences of SpokenTIMs. C-terminal His-tag and linker are underlined.

**Table S2.**
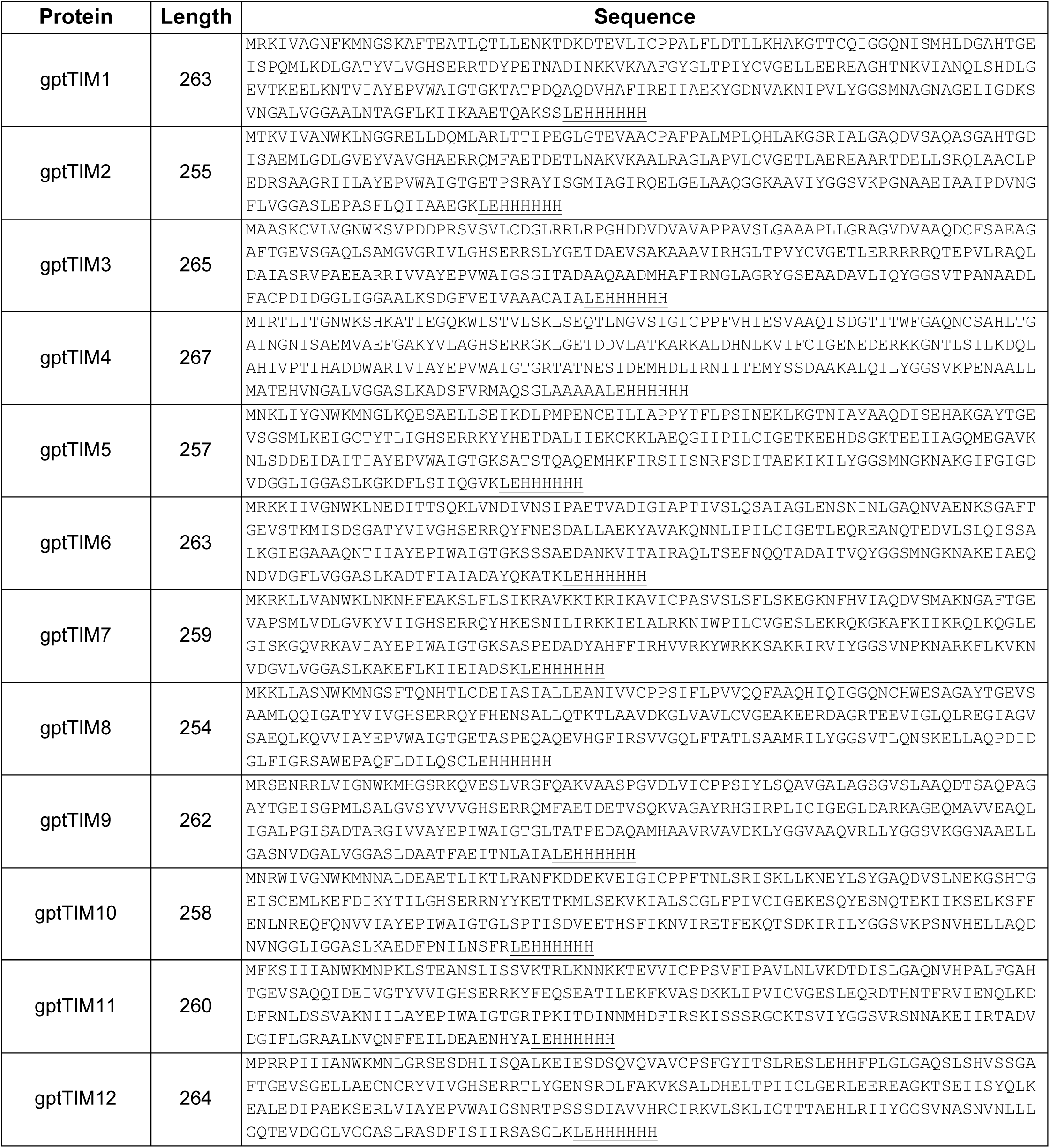
Amino acid sequences of gptTIMs. The C-terminal His-tag and linker are underlined.

**Table S3.** Predicted oligomeric states align with low RMSD values to natural triosephosphate isomerases. Monomeric and dimeric structures of the designs were predicted using ColabFold and aligned to the triosephosphate isomerase of *Clostridium perfringens* (PDB: 4Y8F). All design models superpose to the natural dimer with low RMSD values.

| Protein | RMSD vs. natural monomer TIM<br>(PDB ID: 4Y8F) |  | RMSD vs. natural dimer TIM<br>(PDB ID: 4Y8F) |  |
| --- | --- | --- | --- | --- |
|  | RMSD (Å) | Cα (number) | RMSD (Å) | Cα (number) |
| SpokenTIM1 | 2.04 | 205 | 2.06 | 378 |
| SpokenTIM2 | 1.68 | 189 | 2.25 | 379 |
| SpokenTIM3 | 1.30 | 207 | 1.44 | 391 |
| SpokenTIM4 | 2.13 | 172 | 2.38 | 318 |
| SpokenTIM5 | 1.40 | 200 | 2.12 | 387 |
| SpokenTIM6 | 1.63 | 181 | 2.32 | 382 |
| SpokenTIM7 | 1.59 | 177 | 2.80 | 363 |
| SpokenTIM8 | 1.01 | 197 | 2.14 | 386 |
| SpokenTIM9 | 0.89 | 198 | 1.45 | 377 |
| SpokenTIM10 | 1.31 | 203 | 3.65 | 470 |
| SpokenTIM11 | 1.80 | 188 | 3.01 | 377 |
| SpokenTIM12 | 2.08 | 197 | 2.78 | 383 |
| gptTIM1 | 0.86 | 197 | 1.46 | 448 |
| gptTIM2 | 0.66 | 199 | 0.83 | 406 |
| gptTIM3 | 0.70 | 178 | 1.78 | 424 |
| gptTIM4 | 0.81 | 201 | 3.18 | 470 |
| gptTIM5 | 0.97 | 199 | 1.44 | 458 |
| gptTIM6 | 0.72 | 213 | 1.46 | 471 |
| gptTIM7 | 0.84 | 221 | 0.93 | 420 |
| gptTIM8 | 0.76 | 206 | 1.56 | 454 |
| gptTIM9 | 0.63 | 206 | 0.93 | 445 |
| gptTIM10 | 0.98 | 209 | 1.21 | 460 |
| gptTIM11 | 0.97 | 214 | 1.27 | 428 |
| gptTIM12 | 0.82 | 199 | 1.39 | 456 |

**Table S4.**
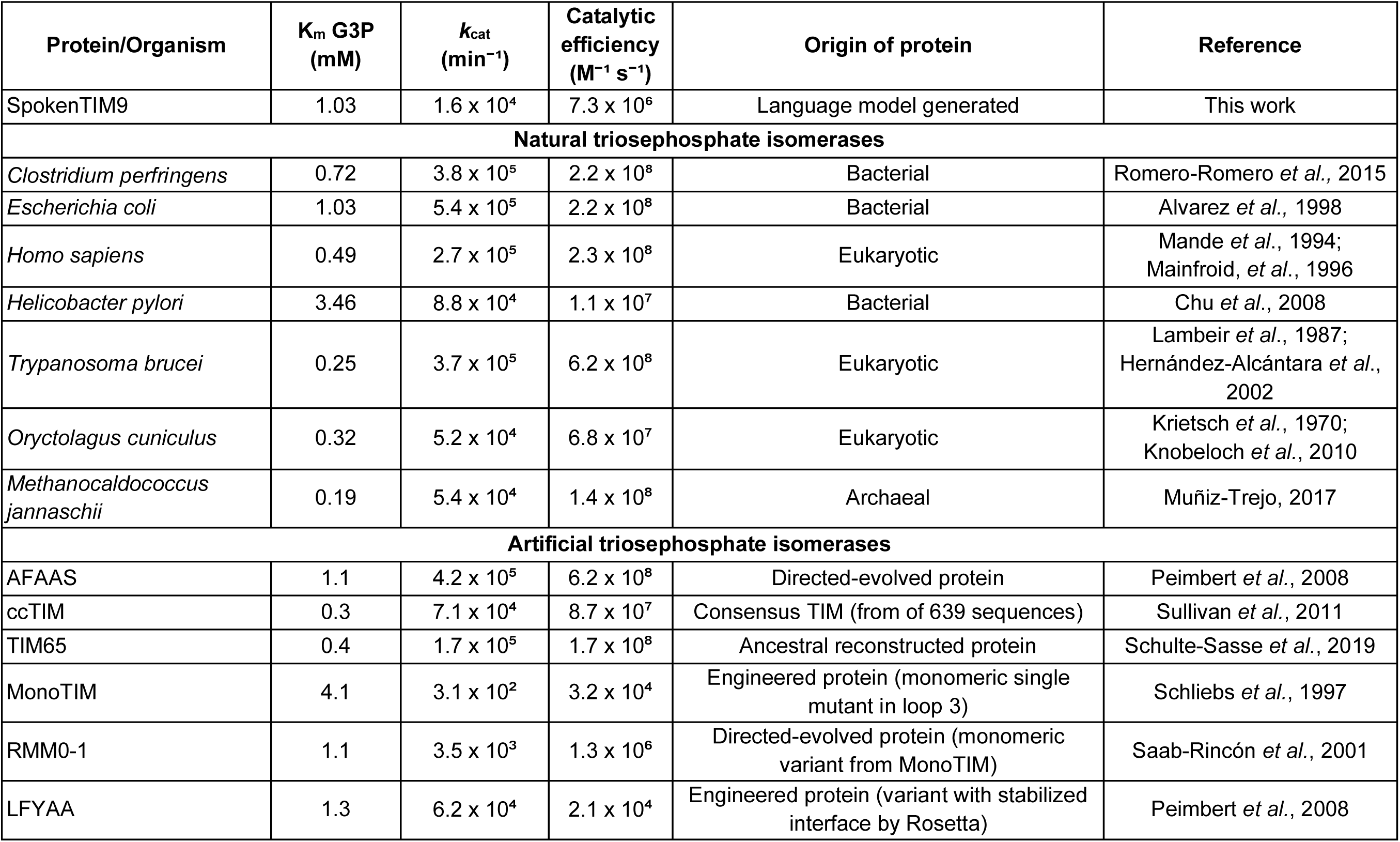
Catalytic parameters reported for natural and artificial TIMs.

## Notes

### Competing Interest Statement

The authors have declared no competing interest.

## References

Alber T, Banner DW, Bloomer AC, Petsko GA, Phillips D, Rivers PS, Wilson IA. 1981. On the three-dimensional structure and catalytic mechanism of triose phosphate isomerase. Philos Trans R Soc Lond B Biol Sci 293: 159–171.

Albery WJ, Knowles JR. 1976. Free-energy profile for the reaction catalyzed by triosephosphate isomerase. Biochemistry 15: 5627–5631.

Alvarez M, Zeelen JPh, Mainfroid V, Rentier-Delrue F, Martial JA, Wyns L, Wierenga RK, Maes D. 1998. Triose-phosphate Isomerase (TIM) of the Psychrophilic BacteriumVibrio marinus. J Biol Chem 273: 2199–2206.

Baek M, DiMaio F, Anishchenko I, Dauparas J, Ovchinnikov S, Lee GR, Wang J, Cong Q, Kinch LN, Schaeffer RD, et al. 2021. Accurate prediction of protein structures and interactions using a three-track neural network. Science 373: 871–876.

Banner DW, Bloomer AC, Petsko GA, Phillips DC, Pogson CI, Wilson IA, Corran PH, Furth AJ, Milman JD, Offord RE, et al. 1975. Structure of chicken muscle triose phosphate isomerase determined crystallographically at 2.5Å resolution: using amino acid sequence data. Nature 255: 609–614.

Bjelic S, Nivón LG, Çelebi-Ölçüm N, Kiss G, Rosewall CF, Lovick HM, Ingalls EL, Gallaher JL, Seetharaman J, Lew S, et al. 2013. Computational Design of Enone-Binding Proteins with Catalytic Activity for the Morita–Baylis–Hillman Reaction. ACS Chem Biol 8: 749–757.

Camacho C, Coulouris G, Avagyan V, Ma N, Papadopoulos J, Bealer K, Madden TL. 2009. BLAST+: architecture and applications. BMC Bioinformatics 10: 421.

Cornish-Bowden A. 2012. Fundamentals of enzyme kinetics. 4., completely revised and greatly enlarged ed. Wiley-Blackwell, Weinheim.

Dauparas J, Anishchenko I, Bennett N, Bai H, Ragotte RJ, Milles LF, Wicky BIM, Courbet A, de Haas RJ, Bethel N, et al. 2022. Robust deep learning–based protein sequence design using ProteinMPNN. Science 378: 49–56.

Dill KA, Ghosh K, Schmit JD. 2011. Physical limits of cells and proteomes. Proc Natl Acad Sci 108: 17876–17882.

Donnelly AE, Murphy GS, Digianantonio KM, Hecht MH. 2018. A de novo enzyme catalyzes a life-sustaining reaction in Escherichia coli. Nat Chem Biol 14: 253–255.

Eddy SR. 2011. Accelerated Profile HMM Searches ed. W.R. Pearson. PLoS Comput Biol 7: e1002195.

Ferruz N, Höcker B. 2022. Controllable protein design with language models. Nat Mach Intell 4: 521–532.

Ferruz N, Michel F, Lobos F, Schmidt S, Höcker B. 2021a. Fuzzle 2.0: Ligand Binding in Natural Protein Building Blocks. Front Mol Biosci 8: 715972.

Ferruz N, Schmidt S, Höcker B. 2021b. ProteinTools: a toolkit to analyze protein structures. Nucleic Acids Res 49: W559–W566.

Ferruz N, Schmidt S, Höcker B. 2022. ProtGPT2 is a deep unsupervised language model for protein design. Nat Commun 13: 4348.

Fleishman SJ, Leaver-Fay A, Corn JE, Strauch E-M, Khare SD, Koga N, Ashworth J, Murphy P, Richter F, Lemmon G, et al. 2011. RosettaScripts: A Scripting Language Interface to the Rosetta Macromolecular Modeling Suite ed. V.N. Uversky. PLoS ONE 6: e20161.

Fox NK, Brenner SE, Chandonia J-M. 2014. SCOPe: Structural Classification of Proteins— extended, integrating SCOP and ASTRAL data and classification of new structures. Nucleic Acids Res 42: D304–D309.

Go MK, Koudelka A, Amyes TL, Richard JP. 2010. Role of Lys-12 in Catalysis by Triosephosphate Isomerase: A Two-Part Substrate Approach. Biochemistry 49: 5377–5389.

Goyal VD, Yadav P, Kumar A, Ghosh B, Makde RD. 2014. Crystallization and preliminary X-ray crystallographic analysis of an artificial molten-globular-like triosephosphate isomerase protein of mixed phylogenetic origin. Acta Crystallogr Sect F Struct Biol Commun 70: 1521–1525.

Herud-Sikimić O, Stiel AC, Kolb M, Shanmugaratnam S, Berendzen KW, Feldhaus C, Höcker B, Jürgens G. 2021. A biosensor for the direct visualization of auxin. Nature 592: 768–772.

Huang P-S, Boyken SE, Baker D. 2016. The coming of age of de novo protein design. Nature 537: 320–327.

Jumper J, Evans R, Pritzel A, Green T, Figurnov M, Ronneberger O, Tunyasuvunakool K, Bates R, Žídek A, Potapenko A, et al. 2021. Highly accurate protein structure prediction with AlphaFold. Nature 596: 583–589.

Khakzad H, Igashov I, Schneuing A, Goverde C, Bronstein M, Correia B. 2023. A new age in protein design empowered by deep learning. Cell Syst 14: 925–939.

Knowles JR. 1991. Enzyme catalysis: not different, just better. Nature 350: 121–124.

Korendovych IV, DeGrado WF. 2020. *De novo* protein design, a retrospective. Q Rev Biophys 53: e3.

Korendovych IV, Kulp DW, Wu Y, Cheng H, Roder H, DeGrado WF. 2011. Design of a switchable eliminase. Proc Natl Acad Sci 108: 6823–6827.

Leaver-Fay A, Tyka M, Lewis SM, Lange OF, Thompson J, Jacak R, Kaufman KW, Renfrew PD, Smith CA, Sheffler W, et al. 2011. Rosetta3. In Methods in Enzymology, Vol. 487 of, pp. 545–574, Elsevier https://linkinghub.elsevier.com/retrieve/pii/B9780123812704000196 (Accessed August 20, 2024).

Lechner H, Ferruz N, Höcker B. 2018. Strategies for designing non-natural enzymes and binders. Curr Opin Chem Biol 47: 67–76.

Lin Z, Akin H, Rao R, Hie B, Zhu Z, Lu W, Smetanin N, Verkuil R, Kabeli O, Shmueli Y, et al. 2023. Evolutionary-scale prediction of atomic-level protein structure with a language model. Science 379: 1123–1130.

Madani A, Krause B, Greene ER, Subramanian S, Mohr BP, Holton JM, Olmos JL, Xiong C, Sun ZZ, Socher R, et al. 2023. Large language models generate functional protein sequences across diverse families. Nat Biotechnol 41: 1099–1106.

Mirdita M, Schütze K, Moriwaki Y, Heo L, Ovchinnikov S, Steinegger M. 2022. ColabFold: making protein folding accessible to all. Nat Methods 19: 679–682.

Moroz YS, Dunston TT, Makhlynets OV, Moroz OV, Wu Y, Yoon JH, Olsen AB, McLaughlin JM, Mack KL, Gosavi PM, et al. 2015. New Tricks for Old Proteins: Single Mutations in a Nonenzymatic Protein Give Rise to Various Enzymatic Activities. J Am Chem Soc 137: 14905–14911.

Munsamy G, Illanes-Vicioso R, Funcillo S, Nakou IT, Lindner S, Ayres G, Sheehan LS, Moss S, Eckhard U, Lorenz P, et al. 2024. Conditional language models enable the efficient design of proficient enzymes. http://biorxiv.org/lookup/doi/10.1101/2024.05.03.592223 (Accessed August 19, 2024).

Nagano N, Orengo CA, Thornton JM. 2002. One Fold with Many Functions: The Evolutionary Relationships between TIM Barrel Families Based on their Sequences, Structures and Functions. J Mol Biol 321: 741–765.

Nakamura T, Yamada KD, Tomii K, Katoh K. 2018. Parallelization of MAFFT for large-scale multiple sequence alignments ed. J. Hancock. Bioinformatics 34: 2490–2492.

Olivares-Illana V, Riveros-Rosas H, Cabrera N, Tuena de Gómez-Puyou M, Pérez-Montfort R, Costas M, Gómez-Puyou A. 2017. A guide to the effects of a large portion of the residues of triosephosphate isomerase on catalysis, stability, druggability, and human disease. Proteins Struct Funct Bioinforma 85: 1190–1211.

Orosz F, Oláh J, Ovádi J. 2006. Triosephosphate isomerase deficiency: Facts and doubts. IUBMB Life 58: 703–715.

Pace N, Scholtz JM, Grimsley GR. 2014. Forces stabilizing proteins. FEBS Lett 588: 2177–2184.

Pan X, Kortemme T. 2021. Recent advances in de novo protein design: Principles, methods, and applications. J Biol Chem 296: 100558.

Peimbert M, Domínguez-Ramírez L, Fernández-Velasco DA. 2008. Hydrophobic Repacking of the Dimer Interface of Triosephosphate Isomerase by in Silico Design and Directed Evolution. Biochemistry 47: 5556–5564.

Polizzi NF, DeGrado WF. 2020. A defined structural unit enables de novo design of small-molecule–binding proteins. Science 369: 1227–1233.

Privett HK, Kiss G, Lee TM, Blomberg R, Chica RA, Thomas LM, Hilvert D, Houk KN, Mayo SL. 2012. Iterative approach to computational enzyme design. Proc Natl Acad Sci 109: 3790–3795.

Rajagopalan S, Wang C, Yu K, Kuzin AP, Richter F, Lew S, Miklos AE, Matthews ML, Seetharaman J, Su M, et al. 2014. Design of activated serine–containing catalytic triads with atomic-level accuracy. Nat Chem Biol 10: 386–391.

Rhys GG, Cross JA, Dawson WM, Thompson HF, Shanmugaratnam S, Savery NJ, Dodding MP, Höcker B, Woolfson DN. 2022. De novo designed peptides for cellular delivery and subcellular localisation. Nat Chem Biol 18: 999–1004.

Richard JP. 2012. A Paradigm for Enzyme-Catalyzed Proton Transfer at Carbon: Triosephosphate Isomerase. Biochemistry 51: 2652–2661.

Robertson AD, Murphy KP. 1997. Protein Structure and the Energetics of Protein Stability. Chem Rev 97: 1251–1268.

Romero-Romero S, Costas M, Rodríguez-Romero A, Fernández-Velasco DA. 2015. Reversibility and two state behaviour in the thermal unfolding of oligomeric TIM barrel proteins. Phys Chem Chem Phys 17: 20699–20714.

Romero-Romero S, Garza-Ramos G. 2021. Crystal structure of Triosephosphate Isomerase from Schizosaccharomyces pombe (SpTIM wt). https://www.rcsb.org/structure/7PEJ.

Romero-Romero S, Kordes S, Michel F, Höcker B. 2021. Evolution, folding, and design of TIM barrels and related proteins. Curr Opin Struct Biol 68: 94–104.

Romero-Romero S, Lindner S, Ferruz N. 2023. Exploring the Protein Sequence Space with Global Generative Models. Cold Spring Harb Perspect Biol 15: a041471.

Saab-Rincón G, Juárez VR, Osuna J, Sánchez F, Soberón X. 2001. Different strategies to recover the activity of monomeric triosephosphate isomerase by directed evolution. Protein Eng Des Sel 14: 149–155.

Schliebs W, Thanki N, Eritja R, Wierenga R. 1996. Active site properties of monomeric triosephosphate isomerase (monoTIM) as deduced from mutational and structural studies. Protein Sci 5: 229–239.

Schmirler R, Heinzinger M, Rost B. 2024. Fine-tuning protein language models boosts predictions across diverse tasks. Nat Commun 15: 7407.

Schümperli M, Pellaux R, Panke S. 2007. Chemical and enzymatic routes to dihydroxyacetone phosphate. Appl Microbiol Biotechnol 75: 33–45.

Sesterhenn F, Yang C, Bonet J, Cramer JT, Wen X, Wang Y, Chiang C-I, Abriata LA, Kucharska I, Castoro G, et al. 2020. De novo protein design enables the precise induction of RSV-neutralizing antibodies. Science 368: eaay5051.

Shevchenko G, Sjödin MOD, Malmström D, Wetterhall M, Bergquist J. 2010. Cloud-Point Extraction and Delipidation of Porcine Brain Proteins in Combination with Bottom-Up Mass Spectrometry Approaches for Proteome Analysis. J Proteome Res 9: 3903–3911.

Siegel JB, Zanghellini A, Lovick HM, Kiss G, Lambert AR, St.Clair JL, Gallaher JL, Hilvert D, Gelb MH, Stoddard BL, et al. 2010. Computational Design of an Enzyme Catalyst for a Stereoselective Bimolecular Diels-Alder Reaction. Science 329: 309–313.

Sillitoe I, Bordin N, Dawson N, Waman VP, Ashford P, Scholes HM, Pang CSM, Woodridge L, Rauer C, Sen N, et al. 2021. CATH: increased structural coverage of functional space. Nucleic Acids Res 49: D266–D273.

Soding J, Biegert A, Lupas AN. 2005. The HHpred interactive server for protein homology detection and structure prediction. Nucleic Acids Res 33: W244–W248.

Sullivan BJ, Durani V, Magliery TJ. 2011. Triosephosphate Isomerase by Consensus Design: Dramatic Differences in Physical Properties and Activity of Related Variants. J Mol Biol 413: 195–208.

Tellez LA, Blancas-Mejia LM, Carrillo-Nava E, Mendoza-Hernández G, Cisneros DA, Fernández-Velasco DA. 2008. Thermal Unfolding of Triosephosphate Isomerase from *Entamoeba histolytica* : Dimer Dissociation Leads to Extensive Unfolding. Biochemistry 47: 11665–11673.

Trentham DR, McMurray CH, Pogson CI. 1969. The active chemical state of d -glyceraldehyde 3-phosphate in its reactions with d -glyceraldehyde 3-phosphate dehydrogenase, aldolase and triose phosphate isomerase. Biochem J 114: 19–24.

Van Kempen M, Kim SS, Tumescheit C, Mirdita M, Lee J, Gilchrist CLM, Söding J, Steinegger M. 2023. Fast and accurate protein structure search with Foldseek. Nat Biotechnol. https://www.nature.com/articles/s41587-023-01773-0 (Accessed August 20, 2024).

Vázquez Torres S, Leung PJY, Venkatesh P, Lutz ID, Hink F, Huynh H-H, Becker J, Yeh AH-W, Juergens D, Bennett NR, et al. 2024. De novo design of high-affinity binders of bioactive helical peptides. Nature 626: 435–442.

Ventura S. 2005. Sequence determinants of protein aggregation: tools to increase protein solubility. Microb Cell Factories 4: 11.

Verkuil R, Kabeli O, Du Y, Wicky BIM, Milles LF, Dauparas J, Baker D, Ovchinnikov S, Sercu T, Rives A. 2022. Language models generalize beyond natural proteins. http://biorxiv.org/lookup/doi/10.1101/2022.12.21.521521 (Accessed April 19, 2024).

Villali J, Kern D. 2010. Choreographing an enzyme’s dance. Curr Opin Chem Biol 14: 636–643.

Watson JL, Juergens D, Bennett NR, Trippe BL, Yim J, Eisenach HE, Ahern W, Borst AJ, Ragotte RJ, Milles LF, et al. 2023. De novo design of protein structure and function with RFdiffusion. Nature 620: 1089–1100.

Weiss S, Adolph RS, Schweimer K, DiFonzo A, Meleshin M, Schutkowski M, Steegborn C. 2022. Molecular Mechanism of Sirtuin 1 Modulation by the AROS Protein. Int J Mol Sci 23: 12764.

Zimmermann L, Stephens A, Nam S-Z, Rau D, Kübler J, Lozajic M, Gabler F, Söding J, Lupas AN, Alva V. 2018. A Completely Reimplemented MPI Bioinformatics Toolkit with a New HHpred Server at its Core. J Mol Biol 430: 2237–2243.

## Supplementary References

Alvarez M, Zeelen JP, Mainfroid V, Rentier-Delrue F, Martial JA, Wyns L, Wierenga RK, Maes D. 1998. Triose-phosphate isomerase (TIM) of the psychrophilic bacterium Vibrio marinus. Kinetic and structural properties. J Biol Chem 273: 2199–2206.

Chu C-H, Lai Y-J, Huang H, Sun Y-J. 2008. Kinetic and structural properties of triosephosphate isomerase from Helicobacter pylori. Proteins 71: 396–406.

Goyal VD, Yadav P, Kumar A, Ghosh B, Makde RD. 2014. Crystallization and preliminary X-ray crystallographic analysis of an artificial molten-globular-like triosephosphate isomerase protein of mixed phylogenetic origin. Acta Cryst F 70: 1521–1525.

Hernández-Alcántara G, Garza-Ramos G, Hernández GM, Gómez-Puyou A, Pérez-Montfort R. 2002. Catalysis and stability of triosephosphate isomerase from Trypanosoma brucei with different residues at position 14 of the dimer interface. Characterization of a catalytically competent monomeric enzyme. Biochemistry 41: 4230–4238.

Mainfroid V, Terpstra P, Beauregard M, Frère JM, Mande SC, Hol WG et al. 1996. Three hTIM mutants that provide new insights on why TIM is a dimer. J Mol Biol 257: 441–456.

Mande SC, Mainfroid V, Kalk KH, Goraj K, Martial JA, Hol WG. 1994. Crystal structure of recombinant human triosephosphate isomerase at 2.8 A resolution. Triosephosphate isomerase-related human genetic disorders and comparison with the trypanosomal enzyme. Protein Sci 3: 810–821.

Muñiz-Trejo R. 2020. Mecanismos adaptativos en la evolución de la triosafosfato isomerasa. National Autonomous University of Mexico. https://tesiunam.dgb.unam.mx/.

Peimbert M, Domínguez-Ramírez L, Fernández-Velasco DA. 2008. Hydrophobic repacking of the dimer interface of triosephosphate isomerase by in silico design and directed evolution. Biochemistry 47: 5556–5564.

Romero-Romero S, Costas M, Rodríguez-Romero A, Fernández-Velasco DA. 2015. Reversibility and two state behaviour in the thermal unfolding of oligomeric TIM barrel proteins. Phys chem chem phys 17: 20699–20714.

Saab-Rincón G, Juárez VR, Osuna J, Sánchez F, Soberón X. 2001. Different strategies to recover the activity of monomeric triosephosphate isomerase by directed evolution. Protein engineering 14: 149–155.

Schliebs W, Thanki N, Eritja R, Wierenga RK. 1996. Active site properties of monomeric triosephosphate isomerase (monoTIM) as deduced from mutational and structural studies. Protein Sci 5: 229–239.

Schulte-Sasse M, Pardo-Ávila F, Pulido-Mayoral NO, Vázquez-Lobo A, Costas M, García-Hernández E, et al. 2019. Structural, thermodynamic and catalytic characterization of an ancestral triosephosphate isomerase reveal early evolutionary coupling between monomer association and function. FEBS J 286: 882–900.

Sullivan BJ, Durani V, Magliery TJ. 2011. Triosephosphate isomerase by consensus design: dramatic differences in physical properties and activity of related variants. J Mol Biol 413: 195–208.

Wierenga R K, Hol WG, Misset O, Opperdoes FR. 1984 Preliminary crystallographic studies of triosephosphate isomerase from the blood parasite Trypanosoma brucei brucei. J mol Biol 178: 487–490.

